# Restoration of β-GC trafficking improves the lysosome function in Gaucher’s disease

**DOI:** 10.1101/2022.06.23.497394

**Authors:** Saloni Patel, Dhwani Radhakrishnan, Darpan Kumari, Priyanka Bhansali, Subba Rao Gangi Setty

## Abstract

Lysosomes function as a primary site for catabolism and cellular signaling. These organelles digest a variety of substrates received through endocytosis, secretion and autophagy with the help of resident acid hydrolases. Lysosomal enzymes are folded in the endoplasmic reticulum (ER) and trafficked to lysosomes via Golgi and endocytic route. The inability of hydrolase trafficking due to mutations or mutations in its receptor or cofactor leads to cargo accumulation (storage) in lysosomes, resulting in lysosome storage disorder (LSD). In Gaucher’s disease (GD), the lysosomes accumulate glucosylceramide due to a lack of β-glucocerebrosidase (β-GC) activity that causes lysosome enlargement/dysfunction. We hypothesize that improving the trafficking of mutant β-GC to lysosomes may delay the progression of GD. RNAi screen using high throughput based lysosomal enzyme activity assay followed by reporter trafficking assay utilizing β-GC-mCherry lead to the identification of nine potential phosphatases. Depletion of these phosphatases in HeLa cells enhanced the β-GC activity by increasing the folding and trafficking of Gaucher’s mutants to the lysosomes. Consistently, the lysosomes in primary fibroblasts from GD patients restored their function upon the knockdown of these phosphatases. Thus, these studies provide evidence that altering phosphatome activity possibly delays the GD and forms an alternative therapeutic strategy for this genetic disease.

**Key points:**

- Phosphatome RNAi screen identified both activators and inhibitors of cellular glucocerebrosidase activity
- Depletion of selective phosphatases in HeLa cells improved the folding and trafficking of mutant β-glucocerebrosidase to lysosomes
- Knockdown of selective phosphatases restored the low basal β-glucocerebrosidase activity to that of wild-type in primary cells derived from Gaucher’s disease patients
- Depletion of selective phosphatases displayed variable β-GC activity in neuropathic and non-neuropathic Gaucher’s disease patient cells

## Introduction

Lysosomes are a unique set of organelles that harbor acid hydrolases and are involved in maintaining cellular homeostasis, organelle turnover, intra and extracellular signaling (Ballabio and Bonifacino, 2020; Luzio et al., 2014; Perera and Zoncu, 2016; Platt et al., 2018; Yang and Wang, 2021). These organelles receive the hydrolases from the Golgi via M6PR - dependent and -independent pathways (Saftig and Klumperman, 2009; Schwake et al., 2013; Staudt et al., 2016) and degrade the cargo received from endocytosis, biosynthetic, secretory and autophagy processes (Ballabio and Bonifacino, 2020; Perera and Zoncu, 2016). Apart from the recycling of degraded materials, lysosomes act as a hub for sensing cellular nutrition, inter-organelle communication, wound healing, plasma membrane repair and apoptosis (Ballabio, 2016; Ballabio and Bonifacino, 2020; Huber and Teis, 2016; Perera and Zoncu, 2016). Inherited mutations in the genes of lysosomal hydrolases or their co-activators or receptors result in the loss of lysosome function due to accumulation of respective enzyme’s substrate (storage), observed in lysosome storage disorders (LSDs) (Futerman and van Meer, 2004; Parenti et al., 2015; Platt et al., 2012; Platt et al., 2018; Sun, 2018). Gaucher’s disease (GD) is the most prevalent form of the known 70 LSDs wherein mutations in β-glucocerebrosidase (β-GC, EC 3.2.1.45) enzyme perturbs its trafficking to the lysosomes (Graves et al., 1988). Around 300 mutations in *GBA1* has been reported (Hruska et al., 2008), which leads to the misfolding and degradation of mutant β-GC by ERAD pathway leading to reduced activity in the lysosomes (Ron and Horowitz, 2005; Sawkar et al., 2005; Tan et al., 2014; Wang et al., 2008). The mutations such as N370S and L444P in *GBA1* are the two common forms observed in GD (Hruska et al., 2008). These mutations affect the folding and trafficking of β-GC but not the activity upon their trafficking to the lysosomes resulting in gradual accumulation of the substrate glucosylceramide. This process primarily causes an impairment in lysosome function followed by secondary defects, including atypical activation of signaling pathways that lead to multisystemic dysfunction (Pastores and Hughes, 1993; Platt et al., 2018; Stirnemann et al., 2017). GD has been classified into Type I, II, III and perinatal lethal forms. Additionally, Type II and III are shown to associated with central nervous system (CNS) deficits (Platt et al., 2018; Stirnemann et al., 2017). The currently available treatments such as enzyme replacement therapy (ERT) and substrate reduction therapy (SRT) are acting as therapies for Type I, but not Type II/III, where these reagents do not cross the blood-brain barrier (Cox et al., 2003; Weinreb et al., 2002). Further, a small molecule-based pharmacological chaperone therapy (PCT) is known to improve the folding and trafficking of mutant β-GC (Istaiti et al., 2021), potentially act as a therapy for GD with neuropathic systems and is under clinical trial (Gary et al., 2018; Parenti et al., 2015; Pastores and Hughes, 1993; Platt et al., 2018; Stirnemann et al., 2017).

The modulation of different cellular pathways has been shown to delay GD by restoring the trafficking of mutant β-GC to lysosomes. For example, enhancing the ER calcium levels by blocking ryanodine receptors (Liou et al., 2016; Ong et al., 2010), inhibiting the ERAD-mediated degradation (Ron and Horowitz, 2005) or transcriptional activation of lysosome biogenesis genes through transcription factor (TF) EB (TFEB)/TFE3 (Song et al., 2013) lead to increased folding/trafficking of mutant β-GC and/or lysosomes biogenesis. Interestingly, the alterations in these common cellular pathways may also impact the other forms of LSDs (Song et al., 2013). However, the small molecules modulating the ER and/or lysosome proteostasis are still under clinical trials (Platt et al., 2018). Therefore, extensive studies in identifying the new cellular pathways to fine-tune the folding/trafficking of β-GC to lysosomes are in high priority for curing GD.

The folding and trafficking of lysosomal proteins are a well-tuned process and is tightly regulated by various cellular signaling pathways (Ballabio and Bonifacino, 2020; Kornfeld and Mellman, 1989; Luzio et al., 2014; Saftig and Klumperman, 2009). Phosphorylation is one of the most common post-translational modifications (PTMs) occurring during this signaling. The process of phosphorylation and dephosphorylation is generally controlled by approximately 518 kinases (Manning et al., 2002) and 255 phosphatases (Sacco et al., 2012), respectively and their function in modulating different cellular pathways associated with lysosome function is poorly understood. Phosphatases control the extent and relay of cell signaling by dephosphorylating the biomolecules that are phosphorylated by kinases. Very few phosphatases such as calcineurin (Medina et al., 2015) and MTMR4 have been shown to regulate lysosomal proteostasis through TFEB (Pham et al., 2018), but their role in regulating β-GC folding and trafficking has not been directly elucidated. Similarly, the depletion of RIPK3 has been shown to improve the disease phenotype in a mouse model of GD (Vitner et al., 2014). We hypothesize here that fine-tuning the signaling pathways by modulating the activity of phosphatases may improve the lysosome function or folding/trafficking of mutant β-GC to the lysosomes and can be used to cure/delay the progression of GD. In this study, RNAi screen against the cellular phosphatome (320, including the catalytic and regulatory subunits) identified nine potential phosphatases, which upon depletion in HeLa and primary fibroblasts from GD patients enhanced the lysosome activity by increasing the folding and trafficking of β-GC to lysosomes. Thus, this study provided the potential therapeutic targets for GD and other forms of LSDs.

## Results

β-GC is a lysosomal luminal enzyme that functions at acidic pH and metabolizes the byproduct glucosylceramide into ceramide and glucose (**Fig. 1A**). In GD, β-GC undergoes degradation due to a variety of genetic mutations, which results in the accumulation of glucosylceramide in the lysosomes (Motabar et al., 2012). Moreover, the mutations at N370S and L444P in human *GBA1* are predominant and spread across the Type I, II and III forms of GD (Hruska et al., 2008; Sibille et al., 1993). Our goal is to restore the trafficking of these mutants from ER to Golgi and then to the lysosomes, allowing these β-GC mutants to digest the accumulated glucosylceramide, which delays the disease progression. In this study, we used an RNAi screen to identify the phosphatases that modulate β-GC activity. We then screened these phosphatases for improved trafficking of β-GC mutants to lysosomes. Additionally, the role of phosphatases in regulating the trafficking of lysosome hydrolases, including the mutants of β-GC, to the lysosomes has not yet been investigated. Thus, this approach will help to identify the new druggable targets for GD and other LSDs.

**Figure 1.**
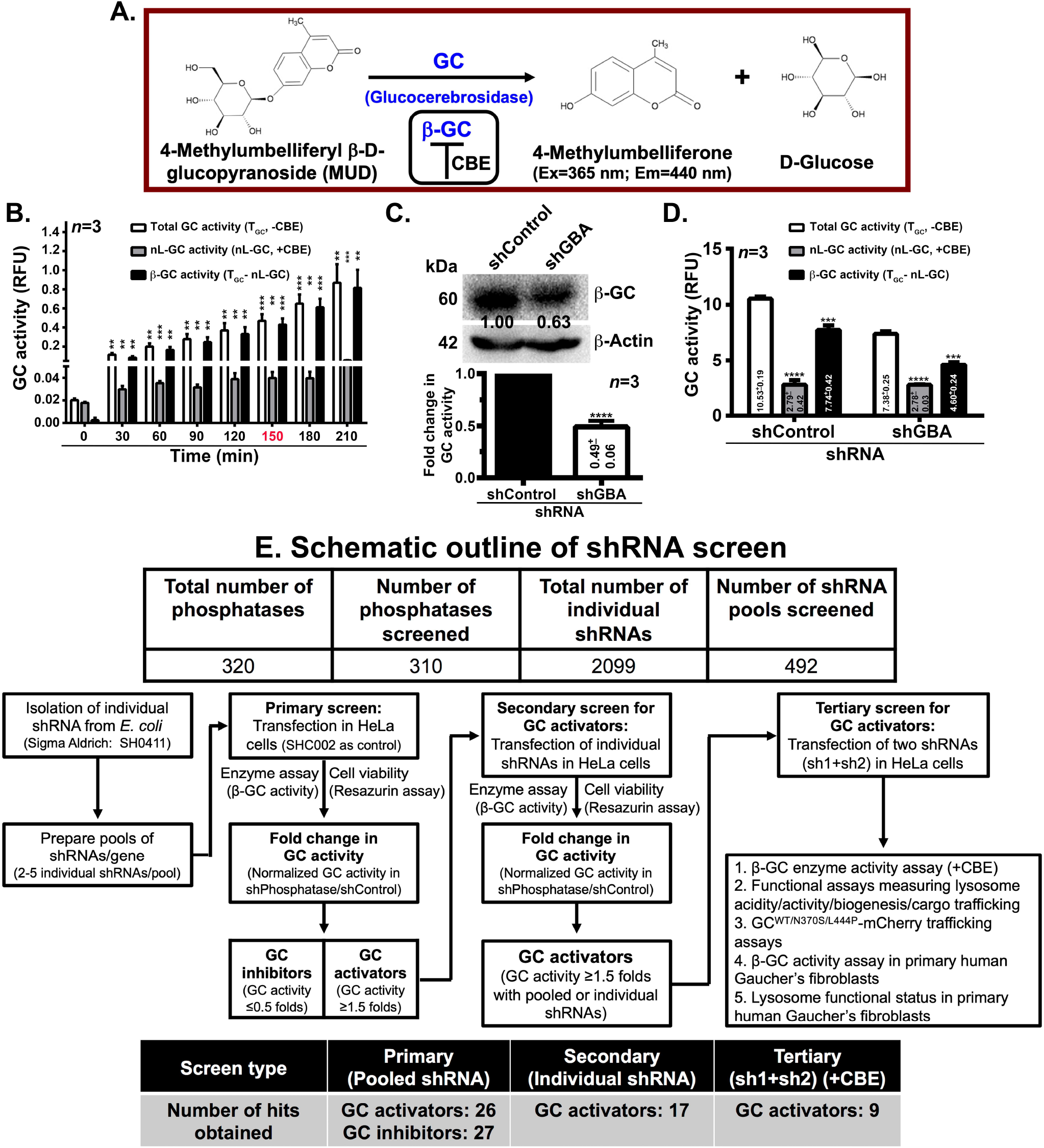
RNAi screen of phosphatome against the intact cell β-GC activity in HeLa cells. (A) Schematic representation of β-GC activity assay. The reaction was carried out in the presence of β-GC inhibitor conduritol B epoxide (CBE). Chemical structures are drawn using ChemDraw JS tool. (B) The plot represents the GC activity (relative fluorescence units) in HeLa cells at different time intervals. The β-GC activity = values of -CBE (total GC, T_GC_) - values of +CBE (nL-GC). The time point 150 min (red color) was chosen for the primary screen. *n*=3. (C) Immunoblotting analysis of GBA knockdown in HeLa cells. The fold change in protein levels is indicated separately. The plot represents the fold change in β-GC activity in GBA-knockdown and control HeLa cells and the value (mean ± s.e.m.) is calculated from D. *n* =3. (D) The plot represents the total GC, nL-GC and β-GC activities in GBA-knockdown and control cells. *n* =3. \*\**p*≤0.01 and \*\*\**p*≤0.001. (E) Schematic outline of RNAi screen of phosphatome against the β-GC activity in HeLa cells. The table represents the total number of phosphatases screened in the assay. The criteria used during primary, secondary and tertiary screening is indicated separately. Flowchart (shown as arrows) also indicate the number of hits obtained at the end of each screening process.

### Screening of phosphatases for modulating the β-GC activity

#### 1. Standardization of β-GC activity assay for the high throughput analysis

To inhibit the activity of phosphatases, we have followed the RNAi approach by using the phosphatome shRNA library (Sigma Aldrich, SH0411). We have performed the transient knockdown of these phosphatases in HeLa cells and measured the β-GC activity. We have standardized the previously published intact cell β-GC activity assay by using 4-methylumbelliferyl β-D-glucopyranoside (MUD, non-fluorescent substrate for both α-glucosidase and β-glucosidase) (Sawkar et al., 2002) amenable for the high-throughput screen (data not shown). Incubation of HeLa cells with MUD in acidic buffer releases a fluorescent product 4-methylumbelliferone (with excitation at 365 nm and emission at 440 nm) and D-glucose by the activity of GC (**Fig. 1A**). Additionally, we have used conduritol B epoxide (CBE), an active site inhibitor of β-GC (Boot et al., 2007), as a control in the assay (**Fig. 1A**).

As reported, cells express non-lysosomal GC (nL-GC, including *GBA2, GBA3* and other enzymes) in addition to β-GC (*GBA1* enzyme) (Boot et al., 2007; van Weely et al., 1993). However, the β-GC functions in an acidic environment, unlike nL-GC (Sawkar et al., 2002). As part of assay standardization to identify the time point and the β-GC specificity, we have incubated the HeLa cells with MUD at variable time points in the presence (+) and absence (-) of CBE (**Fig. 1B**). We have considered the measured GC activity in the presence of CBE as nL-GC or non-specific activity. Plotting the total GC, nL-GC, and β-GC activities separately showed that the β-GC activity values are equivalent to the total GC activity, suggesting that the nL-GC may not show any background activity in the assay condition (**Fig. 1B**). As expected, β-GC activity increases with time and 150 min (2.5 h) was chosen for screening to identify both the up and down regulators of β-GC activity (**Fig. 1B**). For validating the assay, HeLa cells were knockdown for GBA1 using a pool of two shRNAs (**Supplementary Table 1**) and then measured the GC activity in the cells (**Fig. 1C** and **1D**). Consistently, the GBA knockdown cells showed reduced β-GC but not nL-GC activity (**Fig. 1C** and **1D**). Further, the reduced β-GC activity in shGBA cells was almost equivalent to shRNA-mediated gene knockdown efficiency (**Fig. 1C**), indicating the assay is specific to β-GC.

#### 2. Primary screening of phosphatome against β-GC activity

To screen the phosphatases that modulate β-GC activity, we prepared pooled shRNA per gene by mixing DNA of 2 to 5 individual shRNAs, isolated from the glycerol stocks (refer to materials and methods). Note that only a single shRNA was available for few phosphatases in the library and was used in the screen. A total of 492 pools targeting 320 phosphatases (including catalytically active and regulatory subunits, receptors and pseudophosphatases) were prepared manually and transfected into HeLa cells using Lipofectamine 2000 in 24 well plate (**Fig. 1E**). Post transfection and puromycin selection, cells were assayed for cell viability using the resazurin assay followed by β-GC activity assay without CBE. We used non-mammalian targeting shRNA (SHC002, Sigma-Aldrich) as a control throughout the study. As expected, the control shRNA (referred to here as shControl) did not show any toxicity to the cells, and the β-GC activity was nearly equivalent to that of plain cells (data not shown). Fold change in β-GC activity in phosphatome knockdown cells with respect to the control cells was measured and plotted (**Supplementary Fig. 1A** and **1B**). Upon individual knockdown of phosphatases, the β-GC activity in HeLa cells varied from 11.16 to 0.22 folds compared to the control cells (**Supplementary Fig. 1A** and **1B**). In addition, the knockdown cells showed variable cell viability, as shown in the graphs (**Supplementary Fig. 1A** and **1B**). Moreover, the multiple shRNAs pools against the genes ENOPH1, ITPA, PTER, PTPRJ and PTPN1 showed variable GC activity in HeLa cells (data not shown) and therefore excluded for further analysis. Overall, our primary shRNA screen of phosphatome against β-GC activity scored 310 phosphatases, and their activities are ranging from 3.16±0.11 to 0.22±0.05 folds with respective to the control (**Fig. 1E**).

#### 3. Characterization of primary hits into GC activators and inhibitors

The primary screen hits are labeled as GC modulators (not as β-GC modulators) because we did not use CBE in the activity assay. To further screen these modulators, we selected the phosphatases whose knockdown in HeLa cells enhanced the GC activity by 1.5 fold or above compared to the control, defined as GC activators (GC_a_). Similarly, the phosphatases whose knockdown in HeLa cells reduced the activity to 0.5 fold or below compared to the control, defined as GC inhibitors (GC_i_) (**Fig. 1E**). Note that the shPhosphatases showing the GC activity between 0.5 to 1.5 folds with respect to the control were not further examined. Additionally, we did not measure the knockdown efficiency for any shRNA pool during the primary screen. Nevertheless, this analysis identified 26 GC_a_ and 27 GC_i_ phosphatases (**Fig. 2A** and **2B**). We proceeded with the GC_a_ phosphatases to identify the unique phosphatases whose inhibition in activity should enhance the GC activity. Moreover, any small molecular inhibitors available for these phosphatases may substitute the shRNA mediated protein depletion in the cells. In contrast, we predict that the GC_i_ phosphatases may only be useful if we enhance their activity in disease conditions.

**Figure 2.**
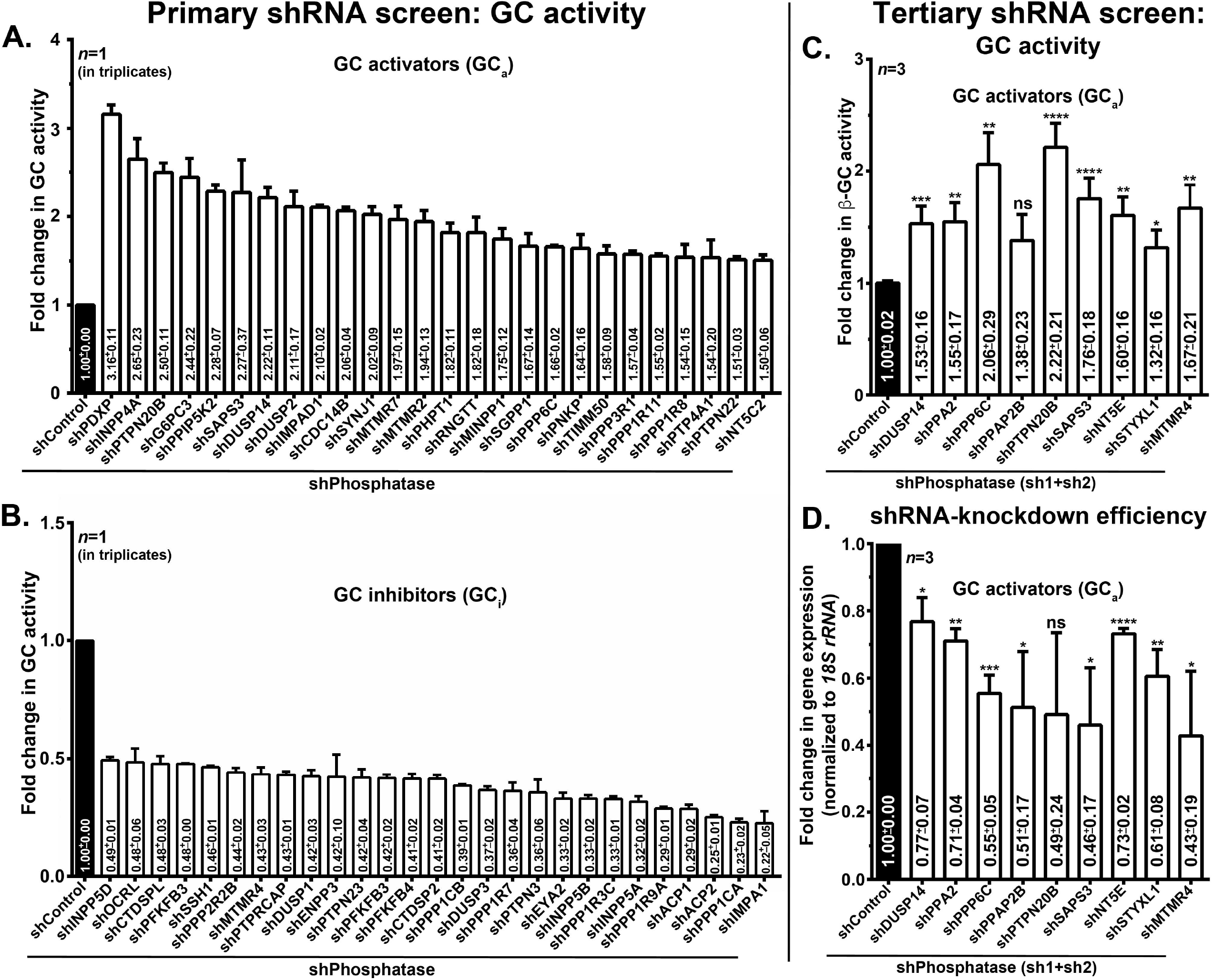
Phosphatome shRNA screen identified novel phosphatases of β-GC activity. (A, B) Plots represent the GC activity of the primary screen hits, labeled as GC activators (GC_a_) and GC inhibitors (GC_i_) separately. Note the GC activity (mean ± s.e.m.) 1.5 fold or above to shControl are scored as GC_a_ phosphatases (A) and 0.5 fold or below to shControl as GC_i_ phosphatases (B). *n*=1 performed in triplicates. (C, D) HeLa cells were knockdown with a mix of shRNAs (sh1+sh2) and shControl and then subjected to β-GC activity (C) and transcript level analysis (D). Fold change in β-GC activity in each phosphatase knockdown compared to shControl was measured and then plotted (mean ± s.e.m.). *n*=3 (each in triplicates). The knockdown efficiency of each gene knockdown in their respective cells in reference to *18S rRNA* was measured and then plotted (mean ± s.e.m.). *n*=3. \**p* ≤0.05, \*\**p*≤0.01, \*\*\**p*≤0.001 and ns=not significant.

#### 4. Secondary and tertiary screening of GC_a_ phosphatases for β-GC activity

To eliminate the false positive hits (due to the off-target effect of shRNAs) of the primary screen, we reanalyzed the 26 GC_a_ phosphatases by repeating the GC activity in HeLa cells that are knockdown with individual shRNAs of each phosphatase (labeled as secondary screen, **Supplementary Fig. 1C**). Interestingly, 8 out of 26 GC_a_ phosphatases retained the GC activity equivalent to the pooled shRNA (**Supplementary Fig. 1C**). Additionally, we have selected 19 phosphatases whose knockdown in HeLa cells showed variable GC activity in different shRNA pools (**Supplementary Fig. 1C**). Repetition of the GC activity in the later set resulted in identifying 9 additional GC_a_ phosphatases (**Supplementary Fig. 1C**). Overall, our secondary screen led to the identification of 17 GC_a_ phosphatases (except shPHPT1, **Supplementary Fig. 1C**), whose knockdown using individual shRNA showed increased GC activity in HeLa cells. Note, two individual shRNAs of PHPT1 showed enhanced while the other two shRNAs decreased the GC activity in HeLa cells compared to control (**Supplementary Fig. 1C**) and, hence, were subjected to tertiary assay.

To determine the specificity of secondary screen hits against β-GC, we selected two best shRNAs (using bioinformatic target sequence analysis) for each phosphatase (**Supplementary Table 1**) that showed enhanced activity in the secondary screen. By using these shRNAs, we performed the GC activity assay in the presence of CBE (labeled as a tertiary screen) and calculated the β-GC activity in the phosphatase knockdown HeLa cells (**Fig. 1E**). In a nutshell, the tertiary screen led to the identification of nine unique phosphatases that are scattered across different phosphatase families. As expected, the knockdown of these phosphatases in HeLa cells enhanced the β-GC activity by 1.6 fold or above with at least one shRNA in the presence of CBE compared to control cells (**Supplementary Fig. 1C**). We have labeled these phosphatases as RNAi hits of β-GC activity. These include DUSP14 and PTPN20B from protein-tyrosine phosphatase superfamily; PPP6C and MTMR4 from protein serine/threonine phosphatase family; PPAP2B, a phospholipid phosphatase; PPA2, a pyrophosphatase; NT5E, a nucleotidase; STYXL1, a pseudophosphatase; and SAPS3, a regulator of phosphatase (**Fig. 1C**). To increase the efficiency of GC_a_ phosphatase knockdown, we combined the two shRNAs selected from the tertiary screen (labeled as shPhosphatase, a mix of sh1+sh2 shown in **Supplementary Fig. 1C** and listed in **Supplementary Table 1**) and was used in the further experiments. As expected, the knockdown of these nine phosphatases using the sh1+sh2 pool in HeLa cells showed a significant (except in shPPAP2B) increase in β-GC activity in HeLa cells compared to the control cells (**Fig. 2C**). However, we observed a variable knockdown (a range from 20% to 60%) of the respective gene upon shRNA (sh1+sh2) transfection in HeLa cells (**Fig. 2D**). Interestingly, the obtained phosphatase knockdown was sufficient to display the enhanced β-GC activity in multiple experiments and improved lysosome function in both HeLa and primary fibroblasts (see below).

### Analyses of GC_a_ phosphatase hits for improved lysosomal function in HeLa cells

Tertiary RNAi screen using β-GC activity assay in HeLa cells identified nine GC_a_ phosphatases (**Fig. 2C**). Interestingly, the role of these phosphatases have not been implicated in the disease pathology linking to GD or elucidated their direct involvement in the lysosome function except MTMR4 (Pham et al., 2018). We tested whether the significant increase in β-GC activity in individual GC_a_ phosphatase knockdown HeLa cells (sh1+sh2) is due to increased lysosome biogenesis/function or enhanced folding/trafficking of β-GC to the lysosomes. To address the earlier point, we measured the following parameters of lysosomes: (1) distribution/position, (2) acidity, (3) proteolytic activity, and (4) status of lysosome biogenesis transcription factors. Upon probing the lysosomes with anti-LAMP-1 antibody, all the GC_a_ phosphatase knockdown HeLa cells showed an enhanced number of lysosomes as compared to shControl cells (**Fig. 3A**). Consistently, we also observed increased dispersal of these lysosomes in the cytosol of all GC_a_ phosphatase knockdown HeLa cells as compared to control cells (quantified as a perinuclear index in **Fig. 3A’**). We observed a minimum dispersion of about 1.21 folds in shMTMR4 and as high as 1.87 folds in shPPAP2B (**Fig. 3A’**).

Lysosomal pH is critical for its function, and recent studies have illustrated a difference in pH between peripheral (less acidic) vs perinuclear (more acidic) localized lysosomes (Johnson et al., 2016). Therefore, we assessed the status of lysosomes by using LysoTracker Red in the GC_a_ phosphatase knockdown and control HeLa cells that are expressing lysosomal protein GFP-LAMP-1 (**Fig. 3B**). Fluorescence microscopy of cells showed the variable number of LysoTracker Red-positive compartments in the GC_a_ phosphatase knockdown and control HeLa cells (**Fig. 3B’**). Interestingly, shDUSP14, shPPA2, shPPP6C, shPTPN20B and shSTYXL1 cells showed an increased number of acidic compartments compared to shControl cells (**Fig. 3B’**). However, we did not observe any change in the LysoTracker Red-positive compartments in the rest of the GC_a_ phosphatase knockdown cells compared to control cells (**Fig. 3B’**). Note that we observed a minor reduction in the number of LysoTracker Red-positive compartments in shPPAP2B cells (**Fig. 3B’**). To know whether the lysosomes in all GC_a_ phosphatase knockdown HeLa cells are acidic, we measured Pearson’s correlation coefficient (*r*) between GFP-LAMP-1 and LysoTracker Red and compared with control cells (**Fig. 3B’’**). Surprisingly, we did not observe any increase in *r* values between GC_a_ phosphatase knockdown and control cells. However, we observed a minor reduction in *r* values in shDUSP14, shPPAP2B, shPTPN20B, shSTYXL1 and shMTMR4 cells (**Fig. 3B’’**). Overall, these studies indicate that the acidity of lysosomes is not altered drastically, even with an increase in lysosome number in certain GC_a_ phosphatases knockdown cells. Moreover, this data suggests that GC_a_ phosphatases may not be involved in maintaining the acidity of lysosomes. Next, we tested the functionality in terms of proteolytic processivity of lysosomes using DQ-Red BSA uptake assay (Marwaha and Sharma, 2017). Upon incubation of cells with DQ-Red BSA, both GC_a_ phosphatase knockdown and control HeLa cells showed DQ-Red fluorescence in LAMP-1-positive compartments (**Fig. 3C**). Quantitative analysis of DQ-Red with LAMP-1 showed no major change in colocalization in all GC_a_ phosphatase knockdown and control HeLa cells (**Fig. 3C’**), suggesting that the knockdown of GC_a_ phosphatases may not affect the processivity of substrates in the lysosomes. Note that we did not observe DQ-Red signal in shPPAP2B cells but restored it upon repetition of the assay in GFP-LAMP-1 transfected shPPA2B cells.

**Figure 3.**
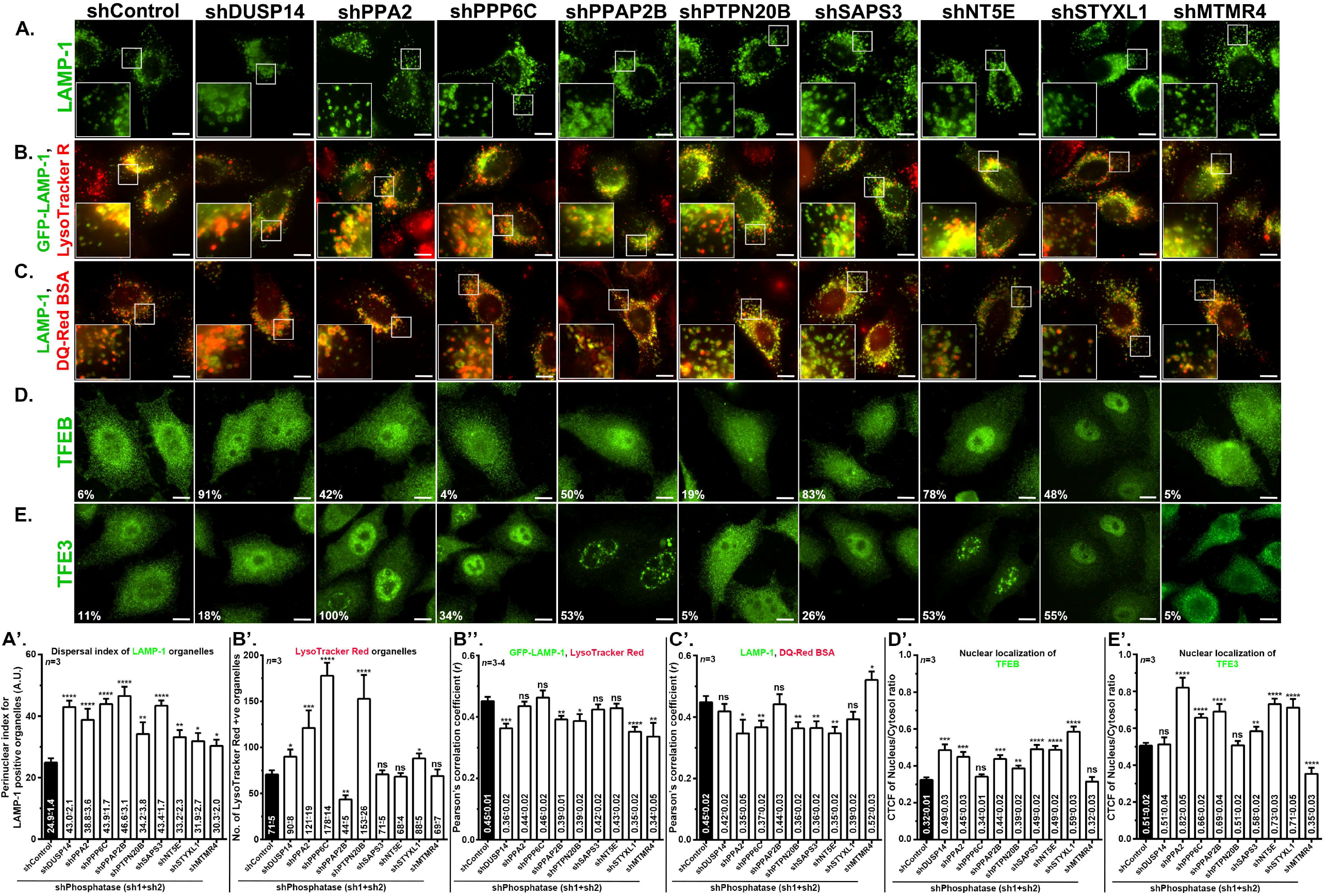
Depletion of GC_a_ phosphatases in HeLa cells alters the lysosome distribution but not biogenesis. (A-E) IFM images of GC_a_ phosphatase knockdown and control HeLa cells. In B-C, cells were incubated with LysoTracker Red (B) or DQ-Red BSA (C). Insets are magnified views of the white boxed areas. Scale bars, 10 μm. Quantification of cells showing the nuclear localization of TFEB or TFE3 is indicated separately (D, E). (A’-E’) Different plots are representing the data shown in A-E. (A’) perinuclear index of LAMP-1-positive organelles in the cells of panel A; (B’) quantification of LysoTracker Red positive compartments in the cells of panel B; (B’’, C’) Pearson’s correlation coefficient (*r*) between LAMP-1 and LysoTracker Red or DQ-Red BSA in the cells of panels B and C respectively; and (D’, E’) quantification of TFEB and TFE3 localization to the nucleus in the cells of panels D and E respectively. \**p* ≤0.05, \*\**p*≤0.01, \*\*\**p*≤0.001 and ns=not significant.

To understand the mechanism behind the increased lysosome numbers or LysoTracker Red-positive compartments or their dispersal in GC_a_ depleted cells, we tested the localization of lysosome biogenesis transcription factors, TFEB and TFE3 (Martina et al., 2016; Settembre et al., 2011) using respective antibodies. Quantification of TFEB localization to the nucleus by immunofluorescence microscopy (IFM) showed TFEB localizes to the nucleus in only 6% of control HeLa cells. We observed a similar percentage of TFEB localization to the nucleus in shPPP6C, shPTPN20B and shMTMR4 cells. In contrast, more than 40% of GC_a_ phosphatases knockdown (shDUSP14, shPPA2, shPPAP2B, shSAP3, shNT5E and shSTYXL1) HeLa cells displayed nuclear TFEB localization. To confirm the results of the above biased-based approach on TFEB localization, we have quantified the ratio of nuclear to cytosol localization of TFEB. Surprisingly, we have observed an increase in the nuclear localization of TFEB in GC_a_ phosphatase knockdown cells compared to control cells except in shPPP6C and shMTMR4 cells (**Fig. 3D’**). Similarly, we observed marginally altered TFE3 localization to the nucleus in the GC_a_ phosphatase knockdown cells compared to control HeLa cells (**Fig. 3E**). The TFE3 was localized to the nucleus in more than 25% of the HeLa cells, especially in shPPA2, PPP6C, shPPAP2B, shSAPS3, shNT5E and shSTYXL1. In other GC_a_ phosphatase knockdown cells, the TFE3 localization to the nucleus was almost similar to that of shcontrol cells (**Fig. 3E**). Quantification of TFE3 localization to the nucleus showed an enhanced nuclear localization in the majority of GC_a_ phosphatase knockdown cells except in shDUSP14, shPTPN20B and shMTMR4 cells compared to control HeLa cells (**Fig. 3E’**). Next, we tested the expression of TFEB/TFE3 driven lysosome biogenesis genes by qRT-PCR. However, the transcript levels of TFEB and TFE3 or their downstream genes did not alter in GC_a_ phosphatase knockdown cells compared to control cells except in shSAPS3 cells (**Supplementary Fig. 2A** and **2B**). In contrast, the levels of TFEB/TFE3 regulated lysosome biogenesis genes moderately reduced in shMTMR4 cells (**Supplementary Fig. 2B**). These results suggest that the knockdown of the GC_a_ phosphatases may not regulate the lysosome biogenesis at the transcriptional level, although a marginally enhanced nuclear localization of TFEB and TFE3 was observed in these cells. Based on these results, we hypothesized that the increased functionality of lysosomes in GC_a_ phosphatase knockdown cells might be due to the increased trafficking of lysosomal proteins such as β-GC to the lysosomes except in shSAP3 cells.

### Reporter cells to measure the trafficking of β-GC to the lysosomes

To test our above prediction, we have generated a β-GC reporter in a lentiviral plasmid wherein the *GBA1* was fused with mCherry epitope tag at the C-terminus (GC^WT^-mCherry) (**Fig. 4A**). Similarly, we wanted to monitor the trafficking of β-GC having GD patient mutations such as N370S and L444P to lysosomes. For this purpose, we generated the GC^N370S^-mCherry and GC^L444P^-mCherry lentiviral reporter constructs through site-directed mutagenesis. Note, N409S and L483P corresponds to N370S and L444P mutations respectively in *GBA1* gene having a leader sequence of 39 aa at the N-terminus (Hruska et al., 2008). Further, we observed an additional mutation at N156D in the GC^N370S^-mCherry construct (**Fig. 4A**). To test the functionality of these reporters, we have generated the lentivirus-mediated stable HeLa cell lines, and they showed an expression of these reporters. Further, we have used a non-clonal HeLa cell population to study the β-GC trafficking to lysosome. Imaging studies revealed that HeLa cells expressing GC^WT^-mCherry localized majorly as punctate structures that are colocalized with lysosomal protein LAMP-1, and a cohort of the reporter observed as reticular pattern positive for calnexin (represents ER) (**Fig. 4B**). In contrast, both GC^N370S^-mCherry and GC^L444P^-mCherry mutants localized majorly to ER and displayed no or minimal localization to lysosomes (**Fig. 4B**). To test the trafficking of GC^N370S/L444P^-mCherry reporters to lysosomes, we have treated the cells with proteasomal inhibitor MG132 (similar to the studies in (Mu et al., 2008)), and these reporters showed localization to LAMP-1-positive lysosomes (**Fig. 4B**). These studies illustrate that the GC-mCherry encoding reporter HeLa cell lines are functional and can be used for testing the β-GC trafficking to the lysosomes. To test whether the stably expressing GC-mCherry reporter cell lines possess any altered β-GC activity in lysosomes, we measured the activity using MUD in the presence of CBE. Compared to empty mCherry expressing cells, GC^WT^-mCherry expressing cells showed increased β-GC activity (1.33±0.16 folds) and decreased in GC^N370S/L444P^-mCherry expressing cells (0.64±0.09 folds in GC^N370S^-mCherry cells, 0.60±0.05 folds in GC^L444P^-mCherry) (**Fig. 4C**). These results suggest that the overexpression of GD mutants of *GBA1* alters the lysosome enzyme activity.

**Figure 4.**
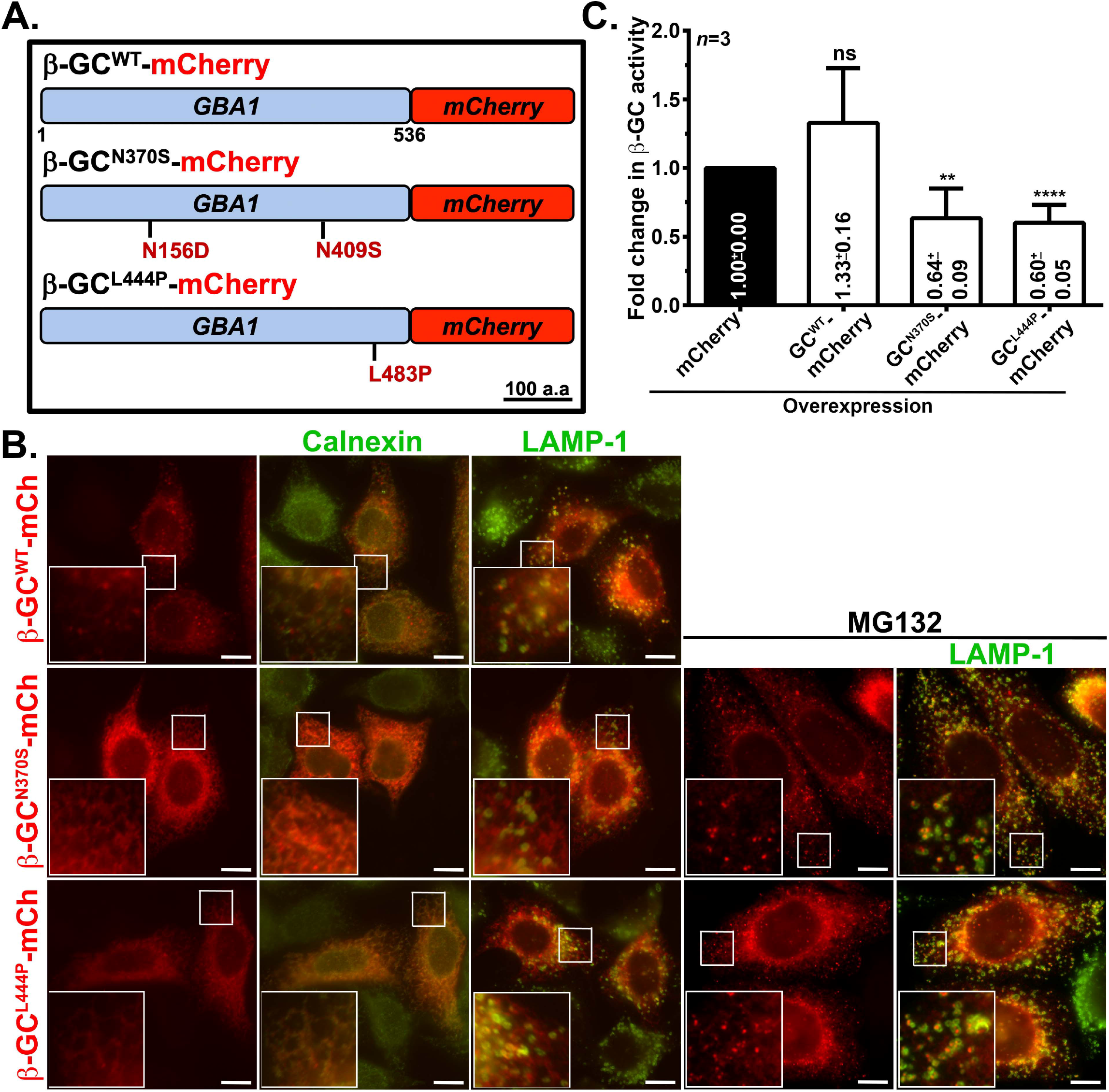
HeLa cells expressing GD mutations encoding GC-mCherry constructs displayed reduced β-GC activity and are targeted to lysosomes upon treatment with MG132. (A) Schematic representation of β-GC^WT/N370S/L444P^-mCherry constructs used in this study. Scale, 100 a.a. Note the presence of additional mutation N156D in β-GC^N370S^-mCherry construct. N409S and L483P are corresponds to N370S and L444P respectively in the isoform-1 of *GBA* gene having a leader sequence of 39 aa at the N-terminus. (B) IFM analysis of stable cells expressing the β-GC^WT/N370S/L444P^-mCherry constructs. Insets are magnified views of the white boxed areas. Scale bars, 10 μm. (C) Plot representing the β-GC activity in HeLa cells stably expressing β-GC-mCherry reporters. An empty vector expressing mCherry is used as a control. Fold change in β-GC activity with respective to the control is indicated (mean ± s.e.m.). *n* =3. \*\**p*≤0.01, \*\*\**p*≤0.001 and ns=not significant.

### Effect of GC_a_ phosphatase knockdown on the folding and trafficking of GC and GD mutants to the lysosome

To test whether the knockdown of GC_a_ phosphatases improves the trafficking of lysosomal hydrolase, β-GC to the lysosomes, we performed the knockdown of GC_a_ phosphatases in reporter cell lines stably expressing GC^WT/N370S/L444P^-mCherry. IFM of these cells revealed that GC^WT^-mCherry localized to LAMP-1-positive lysosomes in all GC_a_ phosphatase knockdown cells, similar to shControl (**Fig. 5A**). The measurement of β-GC^WT^-mCherry puncta and their percentage colocalization with lysosomes in GC_a_ phosphatase knockdown HeLa cells was significantly unaffected compared to control cells (see **Fig. 5D** and **5E**). Interestingly, both GC^N370S^-mCherry and GC^L444P^-mCherry reporters majorly localized to ER in shControl cells and appeared as puncta colocalized with LAMP-1 in all GC_a_ phosphatase knockdown cells (**Fig. 5B** and **5C**). The number of β-GC^N370S/L444P^-mCherry puncta was significantly enhanced in all GC_a_ phosphatase knockdown HeLa cells compared to control cells (see **Fig. 5D and 5E**). As expected, the β-GC^N370S/L444P^-mCherry puncta were colocalized with lysosomes in all phosphatase knockdown and control cells (**Fig. 5D** and **5E**). These studies showed that the knockdown of GC_a_ phosphatases enhanced the trafficking of Gaucher’s mutant reporters to the lysosomes.

**Figure 5.**
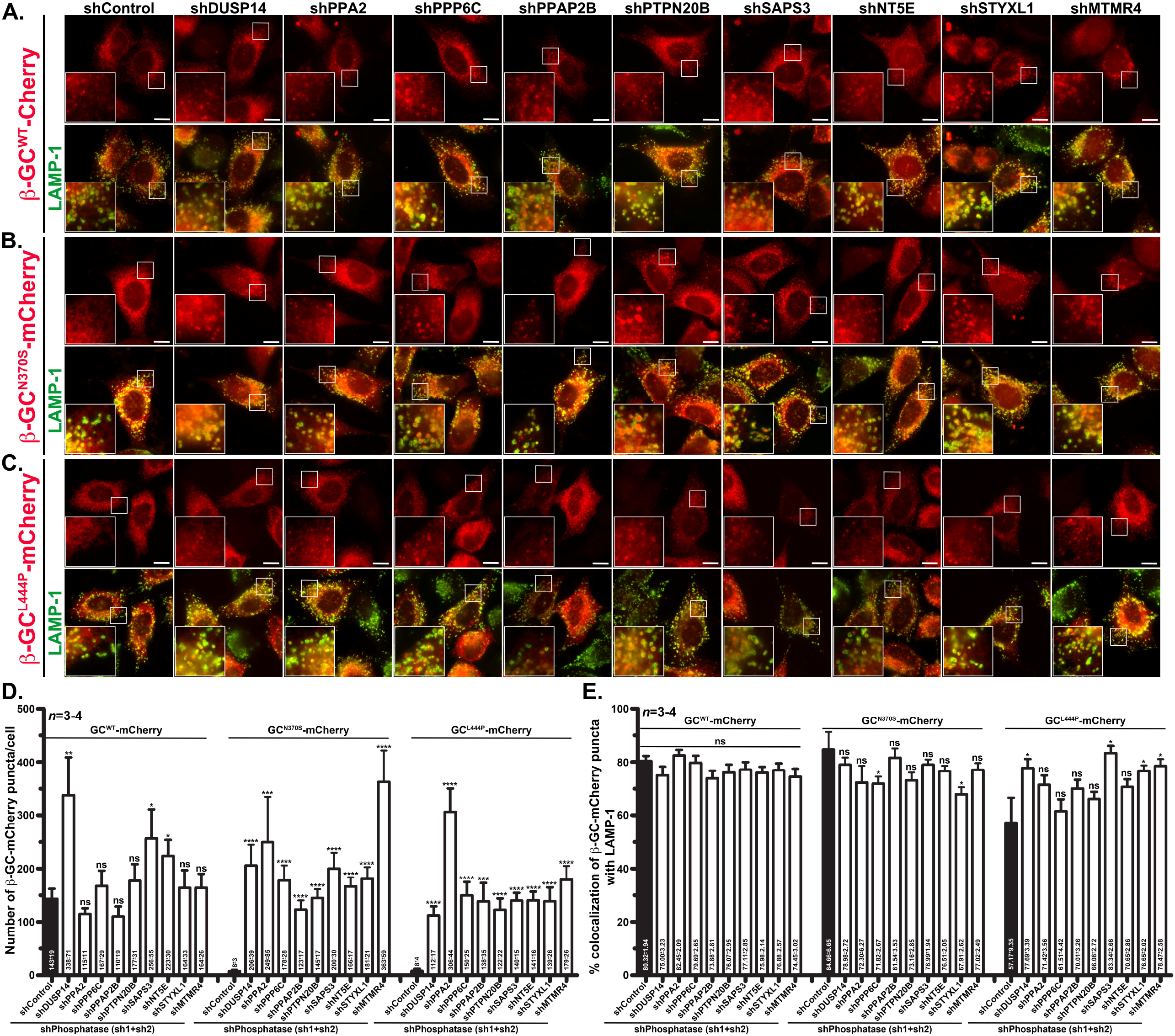
Depletion of GC_a_ phosphatases in HeLa stable cells expressing β-GC^N370S/L444P^-mCherry reporters enhanced their trafficking to lysosomes. (A-C) IFM analyses of HeLa cells expressing β-GC^WT/N370S/L444P^-mCherry reporters. Cells were knockdown for respective phosphatases along with control and then fixed, stained and imaged. Insets are magnified views of the white boxed areas. Scale bars, 10 μm. (D, E) Plots representing the number of β-GC-mCherry puncta/cell (D) or the percentage colocalization of β-GC-mCherry with LAMP-1 (E) in GC_a_ phosphatase-knockdown and control HeLa cells. The average number of puncta or % colocalization of β-GC-mCherry with LAMP-1 (mean±s.e.m.) in each phosphatase knockdown and control cells is indicated. *n* =3-4. \**p* ≤0.05, \*\**p*≤0.01, \*\*\**p*≤0.001 and ns=not significant.

To further confirm that the increased trafficking of GC^N370S/L444P^-mCherry to lysosomes is associated with increased β-GC activity in lysosomes, we have carried out a lysosomal enzyme activity assay using MUD in these cells. Surprisingly, the β-GC activity in GC_a_ phosphatase knockdown HeLa cells significantly increased as compared to control cells (as high as 1.12±0.04 to 3.64±0.11 folds in GC^N370S^-mCherry reporter cells and 2.32±0.17 to 6.35±0.20 folds in GC^L444P^-mCherry reporter cells compared to shControl cells) (**Supplementary Fig. 2C**). These results suggest that the knockdown of these GC_a_ phosphatases in HeLa cells improved the trafficking of GD patient mutants of β-GC to lysosomes.

### Effect of GC_a_ phosphatase knockdown in improving the lysosome activity in Type I and II GD patient fibroblasts

To validate the function of GC_a_ phosphatases in GD, we have obtained the primary human fibroblasts derived from GD patients and wild-type cells (from a healthy individual) (Coriell Life Sciences, USA). For covering the broad spectrum of GD, we have used Type I (GC^N370S,84GG^, GC^N370S,V394L^, GC^L444P^) (GC^L444P^ Type(1) and (2) are from two different patients) and Type II (GC^L444P^) pathology associated GD patient fibroblasts in this study (**Fig. 6**). Initial analysis of GD primary cells by IFM showed enlarged and clustered LAMP-1-positive lysosomes compared to wild-type fibroblasts (**Fig. 6A**). Note that the enlargement of lysosomes is assumed to be due to the accumulation of glucosylceramide in the lysosomes of GD patient cells (Garcia-Sanz et al., 2017; Kinghorn et al., 2017). We have analyzed the cells for lysosome acidity and proteolytic processivity by labeling the cells with LysoTracker Red and DQ-Red BSA, respectively. Interestingly, most of the GD primary fibroblasts showed similar lysosome acidity (quantified as *r* values between LAMP-1 and Lysotracker Red) in both Type I and Type II cells, which is equivalent to wild-type primary cells (**Fig. 6A**). In a similar line, the lysosome protease activity was almost similar in Type I/II GD patient fibroblasts compared to wild-type cells (**Fig. 6A**), suggesting that lysosome acidity or its protease activity was not greatly altered in GD patient fibroblasts. Further, we tested the β-GC activity in these primary cells along with wild-type cells (**Fig. 6B**). As expected, we observed significantly reduced β-GC activity in Type I and II cells than wild-type fibroblasts (**Fig. 6B**). Moreover, the primary cells of GC^N370S,84GG^ and GC^L444P(1)^ of Type I and GC^L444P^ of Type II GD showed a substantial defect in β-GC activity (**Fig. 6B**). These results suggest that GD patient fibroblasts display severe defectivity in β-GC activity due to mutations in *GBA1*; however, their lysosome acidity and general protease activity remain unaffected. Thus, we have used these primary cells to test the effect of GC_a_ phosphatase knockdown on β-GC trafficking to lysosomes.

**Figure 6.**
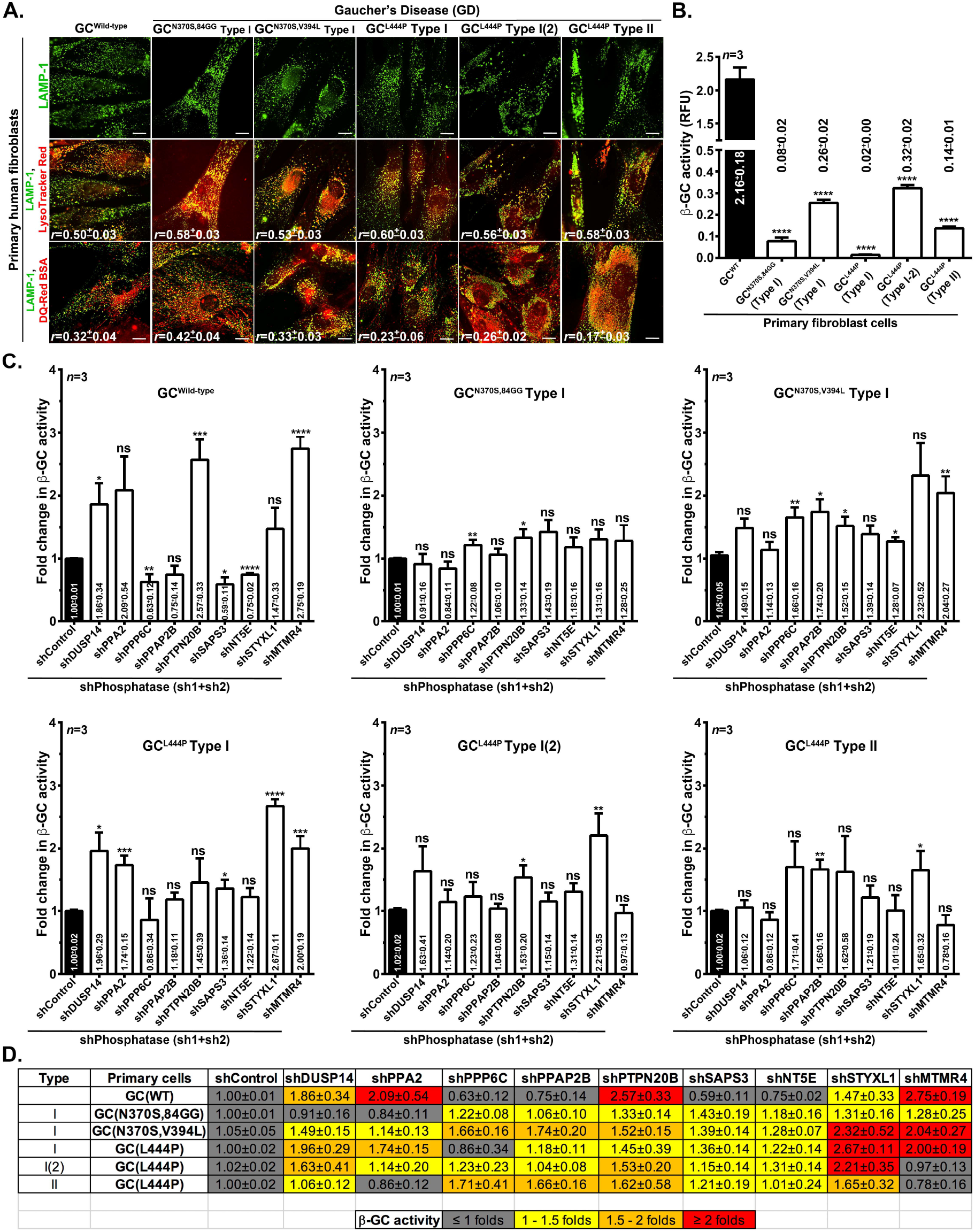
GC_a_ phosphatases knockdown in GD patient fibroblasts improved the β-GC activity. (A) IFM analysis of different types (I and II) of GD and wild-type fibroblasts. Cells were incubated with LysoTracker Red or DQ-BSA Red and then fixed, stained and imaged. The colocalization efficiency (*r*) between LAMP-1 and LysoTracker Red or DQ-Red BSA are indicated in the figures (mean ± s.e.m.). Scale, 10 μm. (B) Plot representing the β-GC activity (relative fluorescence units, mean ± s.e.m.) in GD patient fibroblasts. *n* =3. (C) Different plots representing the β-GC activity in wild-type and Type I and II GD patient fibroblasts post GC_a_ phosphatase knockdown or shControl transduced cells. Fold change in β-GC activity in each GC_a_ phosphatase knockdown cells with respective to the control is indicated (mean ± s.e.m.). *n* =3. \**p* ≤0.05, \*\**p*≤0.01, \*\*\**p*≤0.001 and ns=not significant. (D) Heatmap representing the fold change in β-GC activity as shown in C. Index for the color codes is shown separately.

To analyze the effect of GC_a_ phosphatase knockdown on β-GC activity in wild-type cells, we transduced the primary fibroblasts with lentiviruses encoding respective GC_a_ phosphatase shRNAs (sh1+sh2) or control shRNA. Post 60 h, the cells were assayed for β-GC activity using MUD in the presence of CBE and measured the β-GC activity as fold change in each phosphatase knockdown cells with respect to the control (**Fig. 6C, GC**^**Wild-type**^). In all these experiments, we did not select the primary cells with antibiotic resistance due to their high sensitivity towards the puromycin. Interestingly, the knockdown of phosphatases such as DUSP14, PPA2, PTPN20B, STYXL1 and MTMR4 in wild-type primary fibroblasts enhanced the β-GC activity 1.5 or above folds compared to shControl cells. However, we observed slightly reduced β-GC activity in shPPP6C, shPPAP2B, shSAPS3 and shNT5E cells compared to shControl cells (**Fig. 6C, GC**^**Wild-type**^). Note that the measured β-GC activity in these GC_a_ phosphatase knockdown cells is above the basal level activity shown in **Fig. 6B**. Thus, these studies indicate that the knockdown of certain phosphatases enhances the lysosomal β-GC activity above its basal level. Similar analysis of β-GC activity in Type I GD fibroblasts showed variable phenotypes. Upon knockdown of GC_a_ phosphatases in Type I GD fibroblasts having mutations at N370S and 84GG in β-GC displayed no major change in β-GC activity (between 0.84±0.11 folds to 1.43±0.19 folds), whereas Type I GD fibroblasts having mutations at N370S and V394L in β-GC showed enhanced β-GC activity (between 1.14±0.13 folds to 2.32±0.52 folds) compared to shControl transduced fibroblasts (**Fig. 6C**). Note that the basal level β-GC activity in GC^N370S,84GG^ fibroblasts is significantly lower than in GC^N370S,V394L^ GD fibroblasts (**Fig. 6B**). These studies revealed that the net observed β-GC activity in GC^N370S,84GG^ fibroblasts signifies an improvement in the activity. Interestingly, β-GC activity analysis showed enhanced β-GC activity in the majority of GC_a_ phosphatase knockdown primary fibroblast cells derived from Type I GD patient carrying GC^L444P^ mutation compared to shControl cells (between 1.18±0.11 folds to 2.67±0.11 folds, except shPPP6C in GC^L444P^ Type I and between 1.04±0.08 folds to 2.21±0.35 folds in another patient GC^L444P^ Type I(2)) (**Fig. 6C**). Note that the two different patient cells of GC^L444P^ Type I possess very different basal level β-GC activity (**Fig. 6B**). Consistently, the primary fibroblasts derived from Type II GD patient having L444P mutation displayed an increase in β-GC activity in the majority of GC_a_ phosphatase knockdown cells (except shPPA2 and shMTMR4) compared to shControl cells (between 1.01±0.24 folds to 1.71±0.41 folds) (**Fig. 6C**). Note that these cells possess moderate levels of β-GC activity at the basal level (**Fig. 6B**).

The composite analysis of the β-GC activity in different GD fibroblasts by the heatmap method clearly displayed an increase in β-GC activity by more than 1.01±0.24 folds to 2.67±0.011 folds almost all GC_a_ phosphatase depleted cells compared to control cells (except in few phosphatases labeled as grey) (**Fig. 6D**). In contrast, the knockdown of the majority GC_a_ phosphatases in wild-type cells showed a slight decrease in β-GC activity despite a high level of basal activity (**Fig. 6B** and **6D**). Overall, we observed an improved β-GC activity in most of GD fibroblast cells upon knockdown of GC_a_ phosphatases compared to control cells. Thus, these genes act as potential therapeutic targets in curing GD and other LSDs (required testing in future).

## Discussion

LSDs are known to result from autosomal recessive mutations in the lysosomal hydrolases or their transporters/carrier proteins/cofactors that lead to the accumulation of respective un/partially digested substrate (storage) in the lysosomes (Platt et al., 2018; Sun, 2018). This process not only blocks the recycling of end metabolites but also results in leakage from lysosomes that activates cellular signaling in addition to a change in autophagy flux (Lieberman et al., 2012; Myerowitz et al., 2021; Seranova et al., 2017; Settembre et al., 2008). Moreover, the storage results in a plethora of changes in lysosome homeostasis and affect several tissues/organs, observed as childhood-onset. GD is the most prevalent monogenetic disorder of LSD (Ferreira and Gahl, 2017; Pastores and Hughes, 1993; Platt et al., 2018) and is caused by approximately 300 different familial mutations, grouped into Type I, II and III forms (Grabowski, 1993; Hruska et al., 2008). ERT and SRT have been described as treatments for Type I, but not for neuropathic forms of GD (Futerman and Platt, 2017; Gary et al., 2018). PCT is an alternative and promising strategy for Type II and III GD, which is under clinical trials (Han et al., 2020; Kopytova et al., 2021; Shin and Lim, 2017; Smid et al., 2010). However, these therapies are specific to a type of LSD and mostly act on non-neuropathic forms of the disease. In this study, we have attempted to identify new cellular targets as phosphatases, whose activity modulation may improve the lysosome function even in disease conditions. Moreover, we focused our approach on improving the trafficking of disease mutants of β-GC to lysosomes. These studies illustrated a role for nine new phosphatases, and their depletion improved the lysosomal trafficking of two mutant forms of β-GC and restored the lysosome function in GD condition.

In GD, the mutations in β-GC are subjected to ERAD mediated degradation due to misfolding of the protein in the ER (Tan et al., 2014; Wang and Segatori, 2013; Wang et al., 2008). Interestingly, most of these mutant proteins display an enzyme activity when they are trafficked to lysosomes (**Fig. 6B**) (Lu et al., 2010). Currently, studies such as blocking the calcium release from ryanodine receptor (RYR) (Liou et al., 2016; Ong et al., 2010), enhancing lysosomal biogenesis through TFEB/TFE3 transcription factors (Song et al., 2013), blocking ERAD pathway using MG132 (Ron and Horowitz, 2005) or improving mitochondrial dynamics (Kim et al., 2021) have been shown to alter the trafficking mutant β-GC to lysosomes and restored the lysosome function in GD. However, these targets are broad and may be prone to alter the general physiology of the healthy cells. Nevertheless, the small molecules modulating these pathways are under clinical trials (Platt et al., 2018). However, the approach followed in this study, which is focused on improving the β-GC trafficking through a specific cellular target (phosphatase), may form an alternative strategy to the above methods and has not been explored previously. Moreover, this strategy has several advantages: (1) the assay described here identified several new phosphatases novel to the trafficking of mutant β-GC to the lysosome; (2) the activity of few phosphatases possibly modulated through small inhibitory molecules, those may form potential future drug candidates; (3) the flexibility in altering the phosphatase activity may provide buffering to delay the spectrum of GD mutations and possibly benefit other LSDs.

β-GC functions as a glycolipid hydrolase in lysosomes along with its cofactor saposin C (Liou et al., 2019; Motta et al., 2014; Yoneshige et al., 2015). There are few cellular assays to measure the activity of β-GC and have been used for small molecule screens (Ong et al., 2010; Sawkar et al., 2002). Most in-cell assays utilize an artificial substrate, which will be hydrolysed into a fluorescent molecule by glucocerebroside enzymes (Motabar et al., 2012). In this study, we combined RNAi screen in HeLa cells with a β-GC activity by utilizing MUD as substrate and measured the fluorescence intensity of released 4-methyl umbelliferon using a microplate reader. Additionally, we introduced two conditions for making the assay-specific to β-GC: (1) performed the assay at acidic pH of 4.2, which is equivalent to that of lysosomal pH, and (2) used CBE, an inhibitor specific to β-GC in the reaction buffer as a control (Kuo et al., 2019; Premkumar et al., 2005; Ridley et al., 2013). Both these conditions supported the idea that the measured activity was specific to β-GC but not to nL-GC (**Fig. 1B**). Additionally, we have conducted the study in the form of three different screens (**Fig. 1E**) to reduce the false positive hits. However, the screen has the following pitfalls that may raise the false positives: (1) the efficiency of knockdown between the genes was not measured and possibly varied between the experiments (data not shown); (2) unexpectedly, the multiple shRNA pools for a single gene behaved very differently in the assay (data not shown); and (3) the cell viability was varied between the genes upon phosphatome knockdown in HeLa cells which may alters the measured GC activity (**Supplementary Fig. 1A** and **1B**). We accounted for these possibilities while scoring the hits at the end of each screen; for example, we have considered the gene that showed increased activity in any one of the available shRNA pools (**Supplementary Fig. 1C**). Although we have taken the maximum measures to avoid false positives, we predict that our data might carry few false positives due to unprecedented low resazurin values (cell viability) or the off-target effect of shRNA. The latter was eliminated by scoring the activity using individual shRNAs per gene (during the secondary screen), followed by remeasuring the activity (in the presence of CBE) using a pool of two selective shRNAs (tertiary screen). Additionally, the sensitivity of phosphatase knockdown cells against the puromycin sensitivity was not accounted for in the assay. In contrast, our screen might have missed a few potential positive hits due to the lack of enough shRNA knockdown in the HeLa cells. Nevertheless, our assay conditions overcame these drawbacks and identified both GC_a_ and GC_i_ phosphatases.

Post-screening of genome-wide phosphatome, we predict that the identified 26 GC_a_ phosphatase primary hits may play a key role in improving the lysosome function compared to GC_i_ phosphatases for the following reasons: (1) loss of phosphatase function enhancing GC activity can possibly be mimicked by using small molecules against them in future; (2) these phosphatases may act as inhibitors for lysosomal/GC activity in wild-type cells and would be interesting to study their mechanism of action; and (3) enhancing the GC activity was the primary criteria of this study, which may improve the activity in disease conditions of LSD. Upon screening of 26 primary GC activators further, only four phosphatases retained β-GC activity (considered as hits) upon their knockdown in HeLa cells (**Supplementary Fig. 1C**). At this point, we revisited the genes those having multiple pools of shRNA and selected 19 genes (listed as miscellaneous genes) for the secondary screening. Interestingly, this gene set yielded five new phosphatases as hits after the tertiary shRNA screen (**Supplementary Fig. 1C**). Nonetheless, the identified nine phosphatase hits consistently improved the β-GC activity upon their knockdown in HeLa cells and primary fibroblasts from GD patients. Interestingly, few of these phosphatases have been implicated in lysosome function either directly or indirectly. Studies have shown that the phosphatase MTMR4 has been shown to localize to the late endosomes and autophagosomes and dephosphorylate PI(3)P to PI (Pham et al., 2018). MTMR4 knockdown in A549 cells has been shown to inhibit the nuclear translocation of TFEB following reduced expression of lysosome biogenesis genes upon starvation (Pham et al., 2018). In line with these studies, we did not observe any TFEB and TFE3 translocation to the nucleus upon MTMR4 depletion in HeLa cells (**Fig. 3D, 3E, 3D’** and **3E’**). Moreover, we observed a minor reduction in the expression of lysosome biogenesis genes in the MTMR4 knockdown cells (**Supplementary Fig. 2B**), suggesting the increased β-GC activity and enhanced trafficking of mutant β-GC to lysosomes, is possibly independent of TFEB/TFE3 regulation. Another phosphatase, NT5E (referred as CD73), acts as an ectoenzyme and regulates many lysosome-independent cellular functions (Alcedo et al., 2021). Our studies indicated that this phosphatase might regulate the nuclear translocation of TFEB and TFE3 (**Fig. 3D** and **3E**). However, we did not observe any change in their transcriptional activity (**Supplementary Fig. 2B**), and we predict it may be due to the lower phosphatase knockdown in the cells during these experiments. Interestingly, we have also identified two closely associated phosphatases, PPP6C and SAPS3, to a holoenzyme PP6 in our screen. This heteromeric complex comprises a catalytic (PPP6C), three regulatory (PP6R1/2/3, also called SAPS1/2/3) and three ankyrin repeat-domain containing regulatory subunits (ANKRD28/44/52) (Ohama, 2019). It would be interesting to test whether PPP6C and SAPS3 work together in a PP6 holoenzyme to modulate the β-GC activity or their independent regulation.

The accumulation of glucosylceramide inside the lysosomes is due to mutations in β-GC, which results in the enlargement and dysfunction of lysosomes in GD. This process can be improved by enhancing the folding and trafficking of mutant enzymes or increasing the net lysosomal enzyme activity through lysosome biogenesis. We found that shRNA-mediated depletion of a majority of GC_a_ phosphatase hits in HeLa cells marginally increased the localization of TFEB/TFE3 to the nucleus (**Fig. 3D’** and **3E’**); however, the transcription of lysosome biogenesis genes was not greatly enhanced (**Supplementary Fig. 2B**). Therefore, we predict that the depletion of these phosphatases possibly enhances the folding of mutant

β-GC in ER/Golgi and then the trafficking to lysosomes. As expected, the knockdown of these GC_a_ hits enhanced the folding of endogenous β-GC and the localization of β-GC^N370S/L444P^-mCherry mutants to the lysosomes (positive for LAMP-1) following increased

β-GC activity in HeLa cells (**Fig. 5**). Studies have shown that primary fibroblasts derived from GD patients (Type I, II or III) or the neuronal cells expressing *GBA1* mutants exhibit residual β-GC enzyme activity when compared to those derived from healthy individuals or *GBA1* wild-type respectively (Bae et al., 2015; Ong et al., 2010; Sawkar et al., 2002). Consistently, our studies also observed shallow β-GC activity in GD patient fibroblasts possessing β-GC^N370S^ or β-GC^L444P^ mutation (Type I, II or III) compared with fibroblasts from healthy individuals (**Fig. 6B**). Surprisingly, the lentiviral mediated knockdown of GC_a_ phosphatase hits in GD patient primary fibroblasts carrying different mutations restored the β-GC activity than respective control cells (**Fig. 6D**). Thus, these results provided evidence that the inhibition of GC_a_ phosphatases activity (through gene knockdown) potentially alter the lysosome function through enhanced β-GC trafficking in GD patient cells. Clinical studies indicate that a slight enhancement in enzyme activity may effectively treat GD. For example, infusion of alglucerase enzyme during ERT showed a 1.7 to 9.6 fold increase in β-GC activity in GD patients (Beutler et al., 1995), suggesting a 1.5 to 2 fold increase in activity may be clinically valuable (Sawkar et al., 2002). In support of these studies, we have observed more than one fold GC activity in all our GC_a_ phosphatase knockdown or β-GC^N370S/L444P^-mCherry expressing HeLa cells or GD primary cells (**Figs. 5** and **6**). However, the role of these GC_a_ phosphatases in modulating the lysosome function in other forms of LSD would be interesting to test in future. Overall, our screen identified novel phosphatases, including a pseudophosphatase and subunits of the holoenzyme, whose depletion at the transcript level improved the β-GC activity. Thus, the loss of function of these phosphatases either directly or indirectly alters the cellular pathways, facilitating the trafficking of mutant β-GC to lysosomes. Precisely, identifying the specific substrate or the small inhibitory molecules specific to these phosphatases may provide the mechanism/s underlining the improved lysosome function both in normal cells and primary fibroblasts derived from GD patients.

## Materials and methods

### Reagents and antibodies

All chemicals and tissue culture reagents were purchased either from Sigma-Aldrich (Merck) or ThermoFisher Scientific (Invitrogen). MG132, puromycin, resazurin and 4-methylumbelliferyl β-D-glucopyranoside (MUD) from Sigma-Aldrich (Merck) were purchased. Conduritol B epoxide (CBE) was procured from Tocris Bioscience. LysoTracker Red DND-99 (L7528) and DQ-Red BSA (D12051) were obtained from ThermoFisher Scientific (Invitrogen). Matrigel was purchased from BD Biosciences. Endoglycosidase H (Endo H_f_, P0703L) was obtained from New England BioLabs. The following commercial polyclonal and monoclonal antisera were used (m, mouse; h, human and r, rat proteins): anti-LIMPII (ab165222) was from Abcam; anti-Calnexin (2679), anti-hLAMP-1 (9091) and anti-TFEB (4240) were from Cell Signalling Technology; anti-hLAMP-1 and anti-hLAMP-2 (H4B4) were from Developmental Studies Hybridoma Bank; anti-β-actin (A5441), anti-hGC (G4171), anti-TFE3 (HPA023881) and γ-tubulin (T6557) from Sigma-Aldrich. All secondary antibodies were either from Invitrogen or Jackson Immunoresearch.

### Plasmids and shRNAs

pMD2.G (VSV-G lentiviral envelope vector, 12259) and psPAX2 (lentiviral packaging vector, 12260) were obtained from Addgene. pEGFP-C1-LAMP-1 was obtained as a gift from Dr. Mahak Sharma, IISER Mohali. GC^WT/N370S/L444P^-VSVG-pCDNA3.1+ constructs were a kind gift from Jeffery Kelly, Scripps Institute, USA (Ong et al., 2010). GC^WT/N370S/L444P^-mCherry-pCDNA3.1+ constructs were made by PCR amplifying the gene GC^WT/N370S/L444P^ separately from the above clones, digested and cloned as in-frame N-terminal fusion with mCherry in mCherry-pCDNA3.1+ vector (available in the lab). GC^WT/N370S/L444P^-mCherry-pLVX-puro constructs were made by digesting the GC^WT/N370S/L444P^ -mCherry from their respective pCDNA3.1+ vectors, followed by subcloning into pLVX-puro (Clontech). All constructs were sequenced before use. Note that N409S in GC^N370S^-mCherry and L483P in GC^L444P^-mCherry corresponds to N370S and L444P mutations respectively in the longer isoform-1 of *GBA* (*GBA1*) having a leader sequence of 39 aa at the N-terminus (Hruska et al., 2008). We found an additional N156D mutation in the GC^N370S^-mCherry construct (**Fig. 4A**).

shRNA vectors and phosphatome library: Individual shRNAs against each phosphatase are arrayed as a focused set of phosphatases in human TRC shRNA library, purchased from Sigma-Aldrich, USA (Catalog no. SH0411). All clones are procured as *E. coli* glycerol stocks in a 96 well plate. DNA from each well was isolated using the GenElute HP 96 well plasmid miniprep kit (NA9604). Finally, the DNA was eluted in 150 μl of elution buffer and stored at -20 ^0^C for long term use. The quality and quantity of DNA isolation across the 96 well plate were measured using agarose gel electrophoresis and Nanodrop (ThermoFisher Scientific), respectively. A pool of shRNAs against each phosphatase was prepared by manual mixing of an equal volume of 2-5 shRNAs. Note that several phosphatases possess more than one pool, and we screened as a separate pool in the GC activity assay. pLKO.1-puro Non-mammalian shRNA (referred to here as shControl, SHC002) was used as a control throughout the experiments. In the high throughput screen, we have also included two other additional controls, pLKO.1-puro empty (SHC001) and pLKO.1-puro CMV-Turbo GFP (SHC003).

### Cell culture, transfection and lentiviral transduction

HeLa (ATCC) and HEK293T (ATCC) were maintained in DMEM (Invitrogen) supplemented with 10% FBS (Biowest), 1% L-glutamine (Invitrogen) and 1% penicillin-streptomycin (Pen-Strep, Invitrogen) antibiotics at 10% CO_2_ in a humified cell culture chamber. Cells were transiently transfected with Lipofectamine 2000 (Invitrogen) according to the manufacturer’s protocol with preincubation of cells in OPTI-MEM (Invitrogen) for 30 min. For shRNA screening, HeLa cells were seeded in 24 well plate at an 80% confluency. Post 12 h of plating, cells were transfected with a cocktail containing ∼500 ng of shRNA and

0.75 μl Lipofectamine 2000 in OPTI-MEM. shControl and pLKO.1-puro CMV-Turbo GFP were used as experimental and transfection controls, respectively. Cells were supplemented with a complete medium after 5 h of transfection. Post 48 h, the cells were subjected to two doses of puromycin selection (0.5 μg/ml and 1 μg/ml) for 12 h each with an interval of 12 h between the doses. After selection, the cells were grown in a growth medium without puromycin for 12 h before using it for biochemical GC activity assay or immunofluorescence microscopy. In **Fig. 3B**, cells were transfected with GFP-LAMP-1 post puromycin selection and then plated onto coverslips, fixed, stained and imaged. In **Figs. 4** and **5**, cells were transduced with lentivirus encoding GC^WT^-mCherry, GC^N370S^-mCherry or GC^L444P^-mCherry reporters separately and subjected to puromycin selection to obtain the stable HeLa cells expressing GC reporters. Further, these cells were transiently transfected with shControl or shRNA (mix of sh1 and sh2, sequences are listed in **Supplementary Table 1**) against each phosphatase. Post 48 h, cells were plated on coverslips, fixed, stained and then imaged. Lentiviruses are prepared by following a protocol described previously (Shakya et al., 2018) using HEK293T cells.

Human primary fibroblast cells were maintained in MEM (GIBCO) medium supplemented with 10% FBS (Biowest) and 1% Penicillin-Streptomycin antibiotics (Invitrogen, 100units/ml) at 10% CO2 at 37°C in a humified cell culture chamber. The following primary cells were procured from Coriell Cell Repositories, NJ, USA and used in this study. GC^WT^: apparently health individual from non-fetal tissue, skin origin (GM00498); GC^N370S,84GG^ (Type I): non-neuropathic Gaucher’s disease, Type I, skin origin (GM00372); GC^N370S,V394L^ (Type I): non-neuropathic Gaucher’s disease, Type I, unknown origin (GM01607); GC^L444P^ (Type I): non-neuropathic Gaucher’s disease, Type I, unknown origin (GM10915); GC^L444P^ (Type I-2): non-neuropathic Gaucher’s disease, Type I, skin fetal tissue origin (GM07968); and GC^L444P^ (Type II): acute neuropathic Gaucher’s disease, Type II, unknown origin (GM08760).

### DQ-Red BSA trafficking assay and LysoTracker Red uptake assay

Cells were incubated in a medium containing DQ-Red BSA (10 μg/ml) for 12 hours at 37 ^0^C and then chased in a plain medium for 6 h followed by fixing the cells with 4% paraformaldehyde. Cells were further stained and imaged. For staining LysoTracker Red, cells on the coverslip were incubated in a medium containing LysoTracker Red (100 nM) for 1 h at 37 ^0^C and then fixed with 4% paraformaldehyde. The number of LysoTracker Red-positive puncta/organelles was quantified using Image J. Briefly, the images were converted into binary format followed by applying a common threshold to all images. Further, overlapping objects in the binary images were separated using the Watershed parameter and then quantified the number of puncta by choosing to analyse particle parameters without placing any limit on size and circularity.

### Immunofluorescence microscopy and image analysis

Cells on the coverslip were fixed with 3% formaldehyde and then stained with primary antibodies followed by the respective secondary antibodies described previously (Jani et al., 2015). Immunofluorescence microscopy (IFM) of cells was carried out using a 60X (oil) U Plan super apochromatic objective on an Olympus IX81 motorized inverted fluorescence microscope equipped with a CoolSNAP HQ2 (Photometrics) CCD camera. Images were deconvolved and analyzed with the cellSens Dimension package with the 5D module (Olympus). Analyzed images were assembled using Adobe Photoshop. The colocalization efficiency between two colors was measured by selecting multiple ROIs (region of interests) in the cytosol (excluding perinuclear area) of maximum intensity projection of undeconvolved Z-stack images, followed by estimating the Pearson’s correlation (*r*) value using cellSens Dimension software. The average *r*-value from 20 to 30 cells was calculated and then plotted or represented as mean ± SEM. Nucleus/cytoplasmic ratio of lysosomal transcription factors TFEB and TFE3 was measured by quantifying the corrected total cell fluorescence (CTCF) of their localization to nucleus and cytosol separately. The CTCF was calculated using the below formula and plotted as an averaged ratio between CTCF of the nucleus and CTCF of cytosol from 10 to 20 cells. The mean fluorescence intensity of TFEB/TFE3 localization to the nucleus and cytosol was quantified using Image J. CTCF (in arbitrary units, A.U.) = Integrated density of the cell or nucleus -(area of selected cell or nuclear X mean fluorescence intensity of the background). The distribution of LAMP-1-positive organelles in the cytosol (perinuclear index) was quantified as follows: The fluorescence intensity of LAMP-1 was measured for the entire cell (I_TOTAL_), perinuclear region (the area within 5 μm of the nucleus, I_perinuclear_), and a peripheral region (the area >5 μm from the nucleus, I_peripheral_). Further, the peripheral and perinuclear LAMP-1 intensities were calculated and then normalized using the formula I_>5_ = I_peripheral_/I_total_ − 100 and I_<5_ = I_perinuclear_/I_total_ – 100, respectively. Finally, the perinuclear index was quantified using the formula I_<5_ − I_>5_ × 100. The number of β-GC-mCherry puncta and their percentage colocalization with LAMP-1 were calculated using ImageJ software with ComDet plugin.

### Transcript levels of lysosomal genes by quantitative real-time PCR

Control and phosphatase knockdown cells were subjected to RNA isolation in the presence of β-mercaptoethanol using GeneJET RNA purification kit (ThermoFisher Scientific). The cDNA was synthesized from RNA using a cDNA synthesis kit (Fermantas). The transcripts of lysosome biogenesis genes and their transcription factors were amplified from the cDNA in QuantStudio 6 Flex real-time PCR system (Applied Biosystems). As a control *18S rRNA* or *β-Actin* were amplified along with the lysosomal genes. The C_t_ value and transcript level of each gene relative to the control in phosphatase knockdown cells was quantified and then plotted as fold change to shControl. The primers used in this study are listed in **Supplementary Table 2**.

### Immunoblotting

Cell lysates were prepared in RIPA buffer and then subjected to immunoblotting analysis as described previously (Shakya et al., 2018). Immunoblots were developed with Clarity Western ECL substrate (Bio-Rad) and captured the images in a Bio-Rad Molecular Imager ChemiDoc XRS+ imaging system using Image Lab 4.1 software. Protein band intensities were measured, normalized with loading control, quantified the fold change with respect to control and then indicated in the figure.

### Intact cell β-glucocerebrosidase (GC) activity assay

HeLa cells were seeded in triplicates at 70-80% confluence in 96 well clear flat-bottom black plate (Corning). Cells were subjected to resazurin assay (described below) followed by intact cell lysosomal β-glucosidase activity assay as described previously (Sawkar et al., 2002). Briefly, the cells were washed twice with 1 × PBS post resazurin assay and then incubated with 50 μl of 3 mM MUD (4-methyl umbelliferyl-β-D-glucopyranoside, made in 0.2 M sodium acetate buffer pH 4.2) at 37 °C. The assay was stopped after 150 min (2.5 h) by adding 150 μl of 0.2 M glycine buffer pH 10.8 and measured the fluorescence intensity of liberated 4-methylumbelliferone (excitation at 365 nm and emission at 445 nm) using Tecan multi-mode plate reader. Lastly, the lysosomal enzyme activity (in A.U.) per well was normalized with respective cell viability value. The fold change in GC activity in shPhosphatase with shControl cells was calculated and then plotted. For measuring the activity specific to β-GC, cells in additional triplicate wells were added with 600 μM of CBE (conduritol B epoxide) along with MUD and labeled the obtained activity as +CBE. During the RNAi screen, only one well was added with CBE. Note that the enzymatic reaction was stopped after 60 min during the measurement of β-GC activity in primary human fibroblast cells.

### Cell viability by resazurin assay

HeLa or primary human cells were seeded in triplicates at 70-80% confluence in a 96 well clear flat-bottom black plate (Corning). Resazurin (stock of 10 mg/ml in PBS) was added to the cells to a final concentration of 1 mg/ml in the growth medium and then incubated at 37 ^0^C for 3-4 h. The fluorescence intensity was measured at 530 nm excitation and 590 nm emission using a Tecan multi-mode plate reader (Infinite F200 Pro). The obtained A.U. value was used to normalize the β-GC activity, measured in the same well.

### β-GC-mCherry reporter trafficking assay

HeLa cells stably expressing GC^WT^-mCherry, GC^N370S^-mCherry or GC^L444P^-mCherry were plated directly (in Fig. 4) or post phosphatase knockdown (along with shControl) in either 96 well plate (for β-GC activity) or on coverslips (for immunofluorescence microscopy). In **Fig. 4**, cells on coverslips were subjected to MG132 (10 μM for 4 h) before fixation. Note that the β-GC activity in phosphatase knockdown cells was completed within 48 h of shRNA transfection.

### Statistical analysis

All statistical analyses were carried out using GraphPad Prism 6.0 software. The statistical significance was determined by unpaired Student’s t-test and variance analysis. Ns=not significant, \**p* ≤0.05, \*\**p*≤0.01 and \*\*\**p*≤0.001.

## Acknowledgements

We thank Mahak Sharma for the generous gift of the plasmid GFP-LAMP-1. We thank Jeffery Kelly, Dan Garza, Ting-Wei Mu for sharing the plasmids, protocols, and initial technical support for the GC activity assay. We thank Shruti, Snigdha, Arpita, Ananya, Anup, Madhavi Latha, Sudeshna Nag, Meisam and Krishna for their technical help in handling the genome-wide RNAi library.

## Author contributions

SP performed all the experiments of this study. DR, DK and PB supported the study during multiple steps of shRNA screening, including shRNA mediated knockdown, validation of certain hits by measuring the GC activity and efficiency of gene knockdown, endo H sensitivity assay, and helped in quantifying the immunofluorescence microscopy experiments. SRGS designed, oversaw the entire project, coordinated and discussed the work with co-authors and wrote the manuscript.

## Declarations

### Ethics approval and consent to Participate

Not applicable

### Consent for publication

We give our consent for the publication of identified results described in the manuscript to be published in the journal

### Availability of data and materials

All reagents are available for distribution upon request. The raw data will be shared upon request.

## Supplementary Information

### Supplementary Figures

**Supplementary Figure 1. Primary, secondary and tertiary screening of phosphatome to identify the modulators of GC activity in HeLa cells**. (A, B) Plots represent GC activity (left side) and cell viability (right side) in HeLa cells after phosphatase knockdown. Fold change in GC activity in each phosphatase knockdown cells with respective to the control is plotted. The average resazurin values (relative fluorescence units) are plotted to represent the cell viability. *n*=1 (in triplicates). (C) The table represents the GC activity in the phosphatase knockdown HeLa cells of secondary and tertiary screens. Individual shRNAs per gene was used during the secondary screening, and a pool of two best shRNAs and CBE were used during the tertiary screening. Index for the color codes is shown separately. Each GC activity value is an average of triplicate samples.

**Supplementary Figure 2. Transcript analysis of lysosome biogenesis genes and the transcription factors TFEB and TFE3**. (A, B) qRT-PCR analysis of various lysosome biogenesis transcription factors (A) and genes (B) in GC_a_ phosphatase knockdown and control cell lysates. Fold change (mean ± s.e.m.) in net gene expression with respective to the internal gene control was measured and then plotted. *n*=2. (C) Plot representing the β-GC activity in GC_a_ phosphatase-knockdown HeLa cells stably expressing β-GC^N370S/L444P^-mCherry reporters. Fold change in β-GC activity (mean±s.e.m.) in each phosphatase knockdown cells with respective to the control is indicated. *n* =3. \**p* ≤0.05, \*\**p*≤0.01, \*\*\**p*≤0.001 and ns=not significant.

**Supplementary Table 1**. The table includes the list of selected shRNAs (sh1 and sh2 from the pool) used for each gene knockdown.

**Supplementary Table 2**. The table includes the list of primers used to validate gene knockdown or for transcript-level expression. Both forward and reverse primers, along with the expected band sizes, are indicated in the table.

## References

Alcedo, K. P., Bowser, J. L. and Snider, N. T. (2021). The elegant complexity of mammalian ecto-5’-nucleotidase (CD73). Trends Cell Biol.

Bae, E. J., Yang, N. Y., Lee, C., Lee, H. J., Kim, S., Sardi, S. P. and Lee, S. J. (2015). Loss of glucocerebrosidase 1 activity causes lysosomal dysfunction and alpha-synuclein aggregation. Exp Mol Med 47, e153.

Ballabio, A. (2016). The awesome lysosome. EMBO Mol Med 8, 73–6.

Ballabio, A. and Bonifacino, J. S. (2020). Lysosomes as dynamic regulators of cell and organismal homeostasis. Nat Rev Mol Cell Biol 21, 101–118.

Beutler, E., Kuhl, W. and Vaughan, L. M. (1995). Failure of alglucerase infused into Gaucher disease patients to localize in marrow macrophages. Mol Med 1, 320–4.

Boot, R. G., Verhoek, M., Donker-Koopman, W., Strijland, A., van Marle, J., Overkleeft, H. S., Wennekes, T. and Aerts, J. M. (2007). Identification of the non-lysosomal glucosylceramidase as beta-glucosidase 2. J Biol Chem 282, 1305–12.

Cox, T. M., Aerts, J. M., Andria, G., Beck, M., Belmatoug, N., Bembi, B., Chertkoff, R., Vom Dahl, S., Elstein, D., Erikson, A. et al. (2003). The role of the iminosugar N-butyldeoxynojirimycin (miglustat) in the management of type I (non-neuronopathic) Gaucher disease: a position statement. J Inherit Metab Dis 26, 513–26.

Ferreira, C. R. and Gahl, W. A. (2017). Lysosomal storage diseases. Transl Sci Rare Dis 2, 1–71.

Futerman, A. H. and Platt, F. M. (2017). The metabolism of glucocerebrosides - From 1965 to the present. Mol Genet Metab 120, 22–26.

Futerman, A. H. and van Meer, G. (2004). The cell biology of lysosomal storage disorders. Nat Rev Mol Cell Biol 5, 554–65.

Garcia-Sanz, P., Orgaz, L., Bueno-Gil, G., Espadas, I., Rodriguez-Traver, E., Kulisevsky, J., Gutierrez, A., Davila, J. C., Gonzalez-Polo, R. A., Fuentes, J. M. et al. (2017). N370S-GBA1 mutation causes lysosomal cholesterol accumulation in Parkinson’s disease. Mov Disord 32, 1409–1422.

Gary, S. E., Ryan, E., Steward, A. M. and Sidransky, E. (2018). Recent advances in the diagnosis and management of Gaucher disease. Expert Rev Endocrinol Metab 13, 107–118.

Grabowski, G. A. (1993). Gaucher disease. Enzymology, genetics, and treatment. Adv Hum Genet 21, 377–441.

Graves, P. N., Grabowski, G. A., Eisner, R., Palese, P. and Smith, F. I. (1988). Gaucher disease type 1: cloning and characterization of a cDNA encoding acid beta-glucosidase from an Ashkenazi Jewish patient. DNA 7, 521–8.

Han, T. U., Sam, R. and Sidransky, E. (2020). Small Molecule Chaperones for the Treatment of Gaucher Disease and GBA1-Associated Parkinson Disease. Front Cell Dev Biol 8, 271.

Hruska, K. S., LaMarca, M. E., Scott, C. R. and Sidransky, E. (2008). Gaucher disease: mutation and polymorphism spectrum in the glucocerebrosidase gene (GBA). Hum Mutat 29, 567–83.

Huber, L. A. and Teis, D. (2016). Lysosomal signaling in control of degradation pathways. Curr Opin Cell Biol 39, 8–14.

Istaiti, M., Revel-Vilk, S., Becker-Cohen, M., Dinur, T., Ramaswami, U., Castillo-Garcia, D., Ceron-Rodriguez, M., Chan, A., Rodic, P., Tincheva, R. S. et al. (2021). Upgrading the evidence for the use of ambroxol in Gaucher disease and GBA related Parkinson: Investigator initiated registry based on real life data. Am J Hematol 96, 545–551.

Jani, R. A., Purushothaman, L. K., Rani, S., Bergam, P. and Setty, S. R. (2015). STX13 regulates cargo delivery from recycling endosomes during melanosome biogenesis. J Cell Sci 128, 3263–76.

Johnson, D. E., Ostrowski, P., Jaumouille, V. and Grinstein, S. (2016). The position of lysosomes within the cell determines their luminal pH. J Cell Biol 212, 677–92.

Kim, S., Wong, Y. C., Gao, F. and Krainc, D. (2021). Dysregulation of mitochondria-lysosome contacts by GBA1 dysfunction in dopaminergic neuronal models of Parkinson’s disease. Nat Commun 12, 1807.

Kinghorn, K. J., Asghari, A. M. and Castillo-Quan, J. I. (2017). The emerging role of autophagic-lysosomal dysfunction in Gaucher disease and Parkinson’s disease. Neural Regen Res 12, 380–384.

Kopytova, A. E., Rychkov, G. N., Nikolaev, M. A., Baydakova, G. V., Cheblokov, A. A., Senkevich, K. A., Bogdanova, D. A., Bolshakova, O. I., Miliukhina, I. V., Bezrukikh, V. A. et al. (2021). Ambroxol increases glucocerebrosidase (GCase) activity and restores GCase translocation in primary patient-derived macrophages in Gaucher disease and Parkinsonism. Parkinsonism Relat Disord 84, 112–121.

Kornfeld, S. and Mellman, I. (1989). The biogenesis of lysosomes. Annu Rev Cell Biol 5, 483–525.

Kuo, C. L., Kallemeijn, W. W., Lelieveld, L. T., Mirzaian, M., Zoutendijk, I., Vardi, A., Futerman, A. H., Meijer, A. H., Spaink, H. P., Overkleeft, H. S. et al. (2019). In vivo inactivation of glycosidases by conduritol B epoxide and cyclophellitol as revealed by activity-based protein profiling. FEBS J 286, 584–600.

Lieberman, A. P., Puertollano, R., Raben, N., Slaugenhaupt, S., Walkley, S. U. and Ballabio, A. (2012). Autophagy in lysosomal storage disorders. Autophagy 8, 719–30.

Liou, B., Peng, Y., Li, R., Inskeep, V., Zhang, W., Quinn, B., Dasgupta, N., Blackwood, R., Setchell, K. D., Fleming, S. et al. (2016). Modulating ryanodine receptors with dantrolene attenuates neuronopathic phenotype in Gaucher disease mice. Hum Mol Genet 25, 5126–5141.

Liou, B., Zhang, W., Fannin, V., Quinn, B., Ran, H., Xu, K., Setchell, K. D. R., Witte, D., Grabowski, G. A. and Sun, Y. (2019). Combination of acid beta-glucosidase mutation and Saposin C deficiency in mice reveals Gba1 mutation dependent and tissue-specific disease phenotype. Sci Rep 9, 5571.

Lu, J., Chiang, J., Iyer, R. R., Thompson, E., Kaneski, C. R., Xu, D. S., Yang, C., Chen, M., Hodes, R. J., Lonser, R. R. et al. (2010). Decreased glucocerebrosidase activity in Gaucher disease parallels quantitative enzyme loss due to abnormal interaction with TCP1 and c-Cbl. Proc Natl Acad Sci U S A 107, 21665–70.

Luzio, J. P., Hackmann, Y., Dieckmann, N. M. and Griffiths, G. M. (2014). The biogenesis of lysosomes and lysosome-related organelles. Cold Spring Harb Perspect Biol 6, a016840.

Manning, G., Whyte, D. B., Martinez, R., Hunter, T. and Sudarsanam, S. (2002). The protein kinase complement of the human genome. Science 298, 1912–34.

Martina, J. A., Diab, H. I., Brady, O. A. and Puertollano, R. (2016). TFEB and TFE3 are novel components of the integrated stress response. EMBO J 35, 479–95.

Marwaha, R. and Sharma, M. (2017). DQ-Red BSA Trafficking Assay in Cultured Cells to Assess Cargo Delivery to Lysosomes. Bio Protoc 7.

Medina, D. L., Di Paola, S., Peluso, I., Armani, A., De Stefani, D., Venditti, R., Montefusco, S., Scotto-Rosato, A., Prezioso, C., Forrester, A. et al. (2015). Lysosomal calcium signalling regulates autophagy through calcineurin and TFEB. Nat Cell Biol 17, 288–99.

Motabar, O., Goldin, E., Leister, W., Liu, K., Southall, N., Huang, W., Marugan, J. J., Sidransky, E. and Zheng, W. (2012). A high throughput glucocerebrosidase assay using the natural substrate glucosylceramide. Anal Bioanal Chem 402, 731–9.

Motta, M., Camerini, S., Tatti, M., Casella, M., Torreri, P., Crescenzi, M., Tartaglia, M. and Salvioli, R. (2014). Gaucher disease due to saposin C deficiency is an inherited lysosomal disease caused by rapidly degraded mutant proteins. Hum Mol Genet 23, 5814–26.

Mu, T. W., Ong, D. S., Wang, Y. J., Balch, W. E., Yates, J. R., 3rd, Segatori, L. and Kelly, J. W. (2008). Chemical and biological approaches synergize to ameliorate protein-folding diseases. Cell 134, 769–81.

Myerowitz, R., Puertollano, R. and Raben, N. (2021). Impaired autophagy: The collateral damage of lysosomal storage disorders. EBioMedicine 63, 103166.

Ohama, T. (2019). The multiple functions of protein phosphatase 6. Biochim Biophys Acta Mol Cell Res 1866, 74–82.

Ong, D. S., Mu, T. W., Palmer, A. E. and Kelly, J. W. (2010). Endoplasmic reticulum Ca2+ increases enhance mutant glucocerebrosidase proteostasis. Nat Chem Biol 6, 424–32.

Parenti, G., Andria, G. and Ballabio, A. (2015). Lysosomal storage diseases: from pathophysiology to therapy. Annu Rev Med 66, 471–86.

Pastores, G. M. and Hughes, D. A. (1993). Gaucher Disease. In GeneReviews((R)), (eds M. P. Adam H. H. Ardinger R. A. Pagon S. E. Wallace L. J. H. Bean G. Mirzaa and A. Amemiya). Seattle (WA).

Perera, R. M. and Zoncu, R. (2016). The Lysosome as a Regulatory Hub. Annu Rev Cell Dev Biol 32, 223–253.

Pham, H. Q., Yoshioka, K., Mohri, H., Nakata, H., Aki, S., Ishimaru, K., Takuwa, N. and Takuwa, Y. (2018). MTMR4, a phosphoinositide-specific 3’-phosphatase, regulates TFEB activity and the endocytic and autophagic pathways. Genes Cells.

Platt, F. M., Boland, B. and van der Spoel, A. C. (2012). The cell biology of disease: lysosomal storage disorders: the cellular impact of lysosomal dysfunction. J Cell Biol 199, 723–34.

Platt, F. M., d’Azzo, A., Davidson, B. L., Neufeld, E. F. and Tifft, C. J. (2018). Lysosomal storage diseases. Nat Rev Dis Primers 4, 27.

Premkumar, L., Sawkar, A. R., Boldin-Adamsky, S., Toker, L., Silman, I., Kelly, J. W., Futerman, A. H. and Sussman, J. L. (2005). X-ray structure of human acid-beta-glucosidase covalently bound to conduritol-B-epoxide. Implications for Gaucher disease. J Biol Chem 280, 23815–9.

Ridley, C. M., Thur, K. E., Shanahan, J., Thillaiappan, N. B., Shen, A., Uhl, K., Walden, C. M., Rahim, A. A., Waddington, S. N., Platt, F. M. et al. (2013). beta-Glucosidase 2 (GBA2) activity and imino sugar pharmacology. J Biol Chem 288, 26052–66.

Ron, I. and Horowitz, M. (2005). ER retention and degradation as the molecular basis underlying Gaucher disease heterogeneity. Hum Mol Genet 14, 2387–98.

Sacco, F., Perfetto, L., Castagnoli, L. and Cesareni, G. (2012). The human phosphatase interactome: An intricate family portrait. FEBS Lett 586, 2732–9.

Saftig, P. and Klumperman, J. (2009). Lysosome biogenesis and lysosomal membrane proteins: trafficking meets function. Nat Rev Mol Cell Biol 10, 623–35.

Sawkar, A. R., Adamski-Werner, S. L., Cheng, W. C., Wong, C. H., Beutler, E., Zimmer, K. P. and Kelly, J. W. (2005). Gaucher disease-associated glucocerebrosidases show mutation-dependent chemical chaperoning profiles. Chem Biol 12, 1235–44.

Sawkar, A. R., Cheng, W. C., Beutler, E., Wong, C. H., Balch, W. E. and Kelly, J. W. (2002). Chemical chaperones increase the cellular activity of N370S beta -glucosidase: a therapeutic strategy for Gaucher disease. Proc Natl Acad Sci U S A 99, 15428–33.

Schwake, M., Schroder, B. and Saftig, P. (2013). Lysosomal membrane proteins and their central role in physiology. Traffic 14, 739–48.

Seranova, E., Connolly, K. J., Zatyka, M., Rosenstock, T. R., Barrett, T., Tuxworth, R. I. and Sarkar, S. (2017). Dysregulation of autophagy as a common mechanism in lysosomal storage diseases. Essays Biochem 61, 733–749.

Settembre, C., Di Malta, C., Polito, V. A., Garcia Arencibia, M., Vetrini, F., Erdin, S., Erdin, S. U., Huynh, T., Medina, D., Colella, P. et al. (2011). TFEB links autophagy to lysosomal biogenesis. Science 332, 1429–33.

Settembre, C., Fraldi, A., Jahreiss, L., Spampanato, C., Venturi, C., Medina, D., de Pablo, R., Tacchetti, C., Rubinsztein, D. C. and Ballabio, A. (2008). A block of autophagy in lysosomal storage disorders. Hum Mol Genet 17, 119–29.

Shakya, S., Sharma, P., Bhatt, A. M., Jani, R. A., Delevoye, C. and Setty, S. R. (2018). Rab22A recruits BLOC-1 and BLOC-2 to promote the biogenesis of recycling endosomes. EMBO Rep 19.

Shin, M. H. and Lim, H. S. (2017). Screening methods for identifying pharmacological chaperones. Mol Biosyst 13, 638–647.

Sibille, A., Eng, C. M., Kim, S. J., Pastores, G. and Grabowski, G. A. (1993). Phenotype/genotype correlations in Gaucher disease type I: clinical and therapeutic implications. Am J Hum Genet 52, 1094–101.

Smid, B. E., Aerts, J. M., Boot, R. G., Linthorst, G. E. and Hollak, C. E. (2010). Pharmacological small molecules for the treatment of lysosomal storage disorders. Expert Opin Investig Drugs 19, 1367–79.

Song, W., Wang, F., Savini, M., Ake, A., di Ronza, A., Sardiello, M. and Segatori, L. (2013). TFEB regulates lysosomal proteostasis. Hum Mol Genet 22, 1994–2009.

Staudt, C., Puissant, E. and Boonen, M. (2016). Subcellular Trafficking of Mammalian Lysosomal Proteins: An Extended View. Int J Mol Sci 18.

Stirnemann, J., Belmatoug, N., Camou, F., Serratrice, C., Froissart, R., Caillaud, C., Levade, T., Astudillo, L., Serratrice, J., Brassier, A. et al. (2017). A Review of Gaucher Disease Pathophysiology, Clinical Presentation and Treatments. Int J Mol Sci 18.

Sun, A. (2018). Lysosomal storage disease overview. Ann Transl Med 6, 476.

Tan, Y. L., Genereux, J. C., Pankow, S., Aerts, J. M., Yates, J. R., 3rd and Kelly, J. W. (2014). ERdj3 is an endoplasmic reticulum degradation factor for mutant glucocerebrosidase variants linked to Gaucher’s disease. Chem Biol 21, 967–76.

van Weely, S., Brandsma, M., Strijland, A., Tager, J. M. and Aerts, J. M. (1993). Demonstration of the existence of a second, non-lysosomal glucocerebrosidase that is not deficient in Gaucher disease. Biochim Biophys Acta 1181, 55–62.

Vitner, E. B., Salomon, R., Farfel-Becker, T., Meshcheriakova, A., Ali, M., Klein, A. D., Platt, F. M., Cox, T. M. and Futerman, A. H. (2014). RIPK3 as a potential therapeutic target for Gaucher’s disease. Nat Med 20, 204–8.

Wang, F. and Segatori, L. (2013). Remodeling the proteostasis network to rescue glucocerebrosidase variants by inhibiting ER-associated degradation and enhancing ER folding. PLoS One 8, e61418.

Wang, Q., Li, L. and Ye, Y. (2008). Inhibition of p97-dependent protein degradation by Eeyarestatin I. J Biol Chem 283, 7445–54.

Weinreb, N. J., Charrow, J., Andersson, H. C., Kaplan, P., Kolodny, E. H., Mistry, P., Pastores, G., Rosenbloom, B. E., Scott, C. R., Wappner, R. S. et al. (2002). Effectiveness of enzyme replacement therapy in 1028 patients with type 1 Gaucher disease after 2 to 5 years of treatment: a report from the Gaucher Registry. Am J Med 113, 112–9.

Yang, C. and Wang, X. (2021). Lysosome biogenesis: Regulation and functions. J Cell Biol 220.

Yoneshige, A., Muto, M., Watanabe, T., Hojo, H. and Matsuda, J. (2015). The effects of chemically synthesized saposin C on glucosylceramide-beta-glucosidase. Clin Biochem 48, 1177–80.

