## Supplementary figures and images for "Restoration of β-GC trafficking improves the lysosome function in Gaucher’s disease"

### Supplementary Figure 1

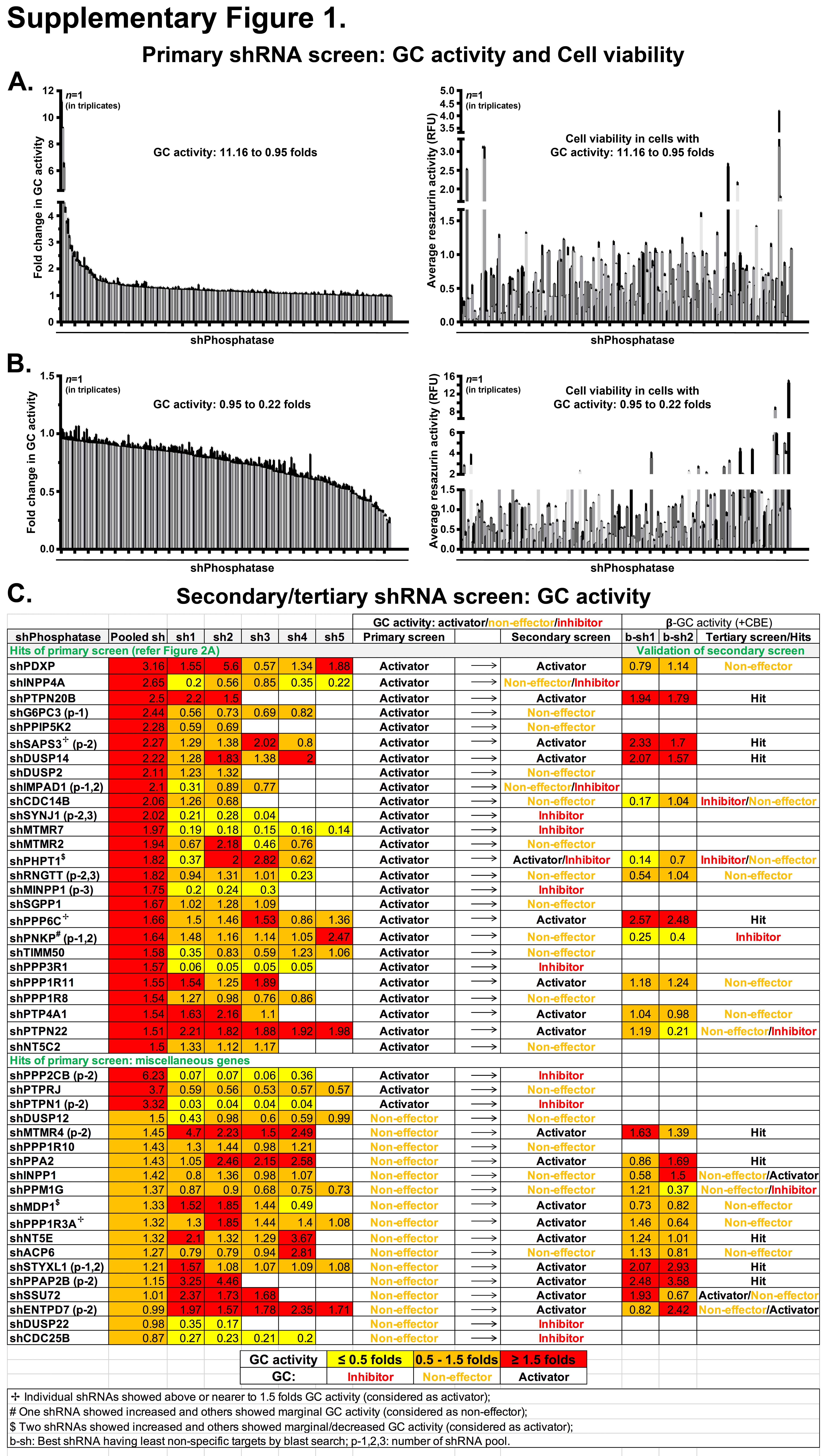

### Supplementary Figure 2

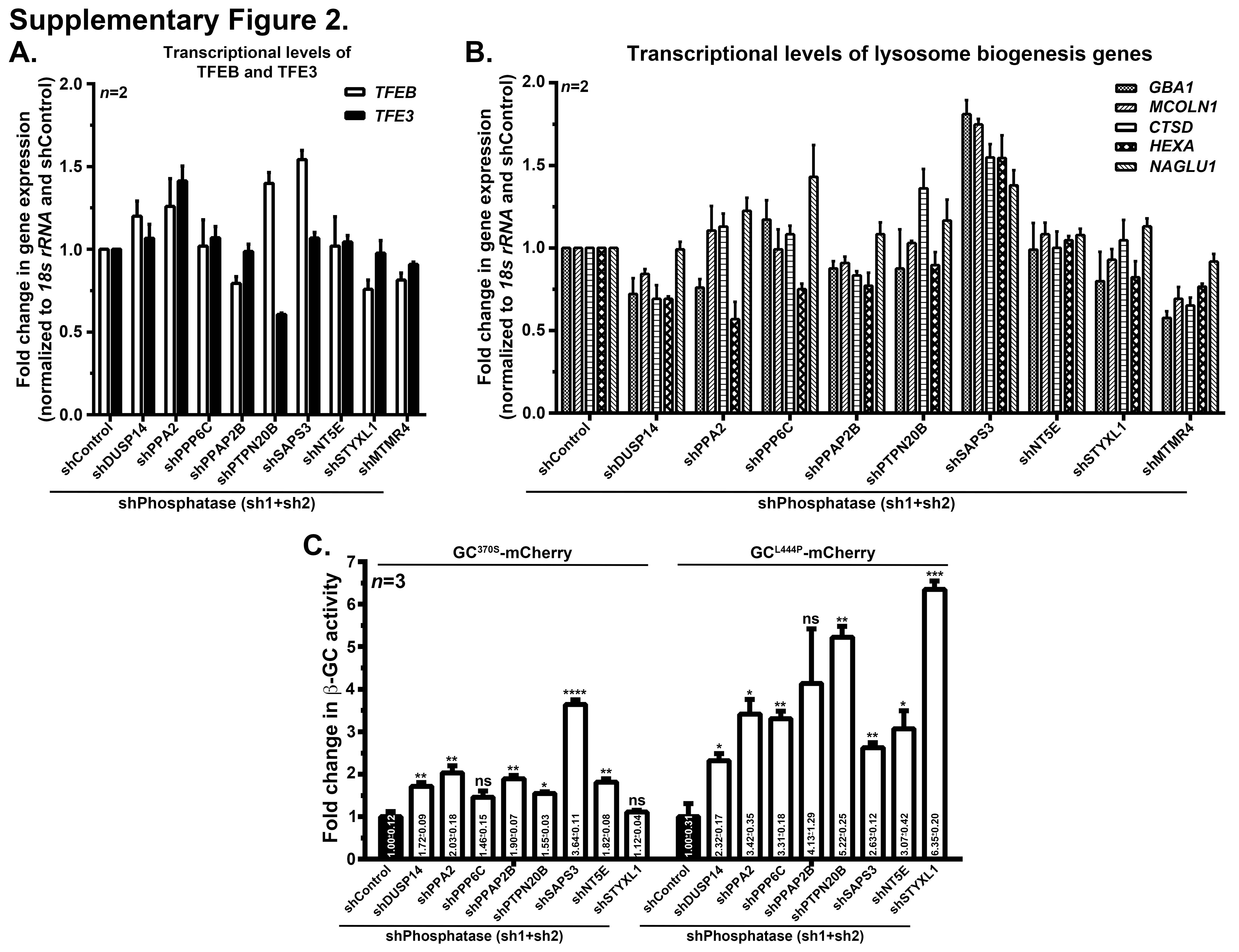
