## Supplementary Table 1 for "Restoration of β-GC trafficking improves the lysosome function in Gaucher’s disease"

**Supplementary Table 1.** List of selected shRNAs used for the gene knockdown

| **Gene** | **shRNA** | **Target sequence: 5’-sequence-3’** |
| --- | --- | --- |
| *GBA1* | sh1  sh2 | CAGTCCCATCATTGTAGACAT  TCTCTGAAGAAGGAATCGGAT |
| *DUSP14* | sh1  sh2 | GCCCAGGATTATATTAGCATT  ATGCTTAGGGAAGGTTGATAA |
| *PPA2* | sh1  sh2 | CCTATGAAGAAAGCACGAAAT  GATCATTAGTTGAATCGGTAT |
| *PPP6C* | sh1  sh2 | GAGTCAAATGTTCAGCCAGTA  CCAAAGTTATTCCGGGCAGTT |
| *PPAP2B* | sh1  sh2 | TCTGACCTCTTCAAGACTAAG  CTGGATGTGCAATTGGTTTAA |
| *PTPN20B* | sh1  sh2 | CCACCCTAACACTTAACATAT  CTTCGGAATTTCCCACATAAT |
| *SAPS3* | sh1  sh2 | CGCAAGAAGAAGATCGACATT  GAATACTTGAAGCCTGGGAAA |
| *NT5E* | sh1  sh2 | CCTCTCAATCATGCCGCTTTA  GCACTGGGAAATCATGAATTT |
| *STYXL1* | sh1  sh2 | GCATAGTAACGAGCAGACCTT  GCTTTACAACATCCTGAATCA |
| *MTMR4* | sh1  sh2 | TAATAGGAATGGGCAGTTATT  GAGCCGTGATATGTTCCAGTT |
