## Supplementary Table 2 for "Restoration of β-GC trafficking improves the lysosome function in Gaucher’s disease"

**Supplementary Table 2.** List of primers used for semi-quantitative PCR and qRT-PCR

| **Gene** | **Forward primer 5’-3’** | **Reverse primer 5’-3’** | **Size (bp)** |
| --- | --- | --- | --- |
| **Control genes** | | | |
| *β-Actin* | CATGTACGTTGCTATCCAGGC | CTCCTTAATGTCACGCACGAT | 250 |
| *18S rRNA* | TTTCGGAACTGAGGCCATGA | GAACCTCCGACTTTCGTTCTTGA | 155 |
| *GAPDH* | AGTCCACTGGCGTCTTCAC | GCTGATGATCTTGAGGCTGT | 155 |
| **Phosphatase genes** | | | |
| *DUSP14* | TCAACTGGCCCCAATTTGAGT | CATCAGGTACGCGATACACAG | 185 |
| *MTMR4* | CCAAGCCAAGGATCTGTTCCC | GCCGGTAGTTAGAGATGGCAA | 143 |
| *NT5E* | GCCTGGGAGCTTACGATTTTG | TAGTGCCCTGGTACTGGTCG | 196 |
| *PPA2* | AAGAAGTTCAAACCGGGTTACC | GCAACGGAAAGGGCTATCAGA | 237 |
| *PPAP2B* | GCTCATCTGCCTCGACCTC | TGATGATCGCGAGGATGGCAA | 200 |
| *PPP6C* | TCCATGGACAGGGTGATTTTGTA | TCCTGATTCCGTTCGATGGT | 325 |
| *PTPN20B* | AGCCTTCAGAAAGTGGCAGT | AAGTGATACTGCTCCTTCGTTT | 997 |
| *SAPS3* | AATGCCACAATTACCGATCAAGA | AAGCTGCTGCACTAATGCACT | 228 |
| *STYXL1* | TCTGTGGACCTGGAGTGTGT | TGGCACGATTTCAATGGGGT | 303 |
| **Lysosome biogenesis genes** | | | |
| *TFEB* | AGTACCTGTCCGAGACCTATG | TGATGTTGAACCTTCGTCTCC | 474 |
| *TFE3* | GCAGGCGATTCAACATTAACG | AAAGTGCAGGTCCAGAAGG | 446 |
| *GBA1* | ACAGGATTGCTTCTACTTCAGG | CGTGTGATTAGCCTGGATGG | 225 |
| *MCOLN1* | CCACATCCAGGAGTGTAAGC | CCAGCCATTGACAAATTCCAG | 226 |
| *CTSD* | ACCTCGTTTGACATCCACTATG | AGGATGCCATCGAACTTGG | 191 |
| *HEXA* | AATGGCGTTAGGGTAAGGGC | ATAGCTCACAGGCAAGGCAG | 122 |
| *NAGLU* | CTTTCAATGAGATGCAGCCAC | TCAGCAAACAGGTCCAGAAC | 220 |
